# Ultra-wide-field, deep, adaptive two-photon microscopy for multi-scale neuronal imaging

**DOI:** 10.1101/2025.06.07.658419

**Authors:** Mengke Yang, Zhen-Qiao Zhou, Song Lang, Hanqing Zheng, Shuai Chen, Tong Li, Eline Stas, Jess Yu, Long Zhang, Zhi Zhang, Volkan Uzungil, Qinying Liu, Yu Huang, Jing Lyu, Yimei Li, Hongbo Jia, Min Li, Xiaojing Li, Jingwei Li, Yuguo Tang, Yan Gong, Simon R. Schultz

## Abstract

Observing the activity patterns of large neural populations throughout the brain is essential for understanding brain function. However, capturing neural interactions across widely distributed brain regions from both superficial and deep cortical layers remains challenging with existing microscopy technologies. Here, we introduce a state-of-the-art two-photon microscopy system, ULTRA, capable of single-cell resolution imaging across an ultra-large field of view (FOV) exceeding 50 mm², enabling deep and very wide field in vivo imaging. To demonstrate its capabilities, we conducted a series of experiments under multiple imaging conditions, successfully visualizing brain structures and neuronal activities spanning a spatial range of over 7 mm from superficial layers to depths of up to 900 μm, while covering a volume of 45.24 mm^3^ in the mouse brain. This versatile imaging platform overcomes traditional spatial constraints, providing a powerful tool for comprehensive exploration of neuronal circuitry over extensive spatial scales with cellular resolution.

## Introduction

Since its inception in the early 1990s, two-photon microscopy (2PM) has become a cornerstone technology for deep-tissue in vivo imaging ^1,2^, offering superior spatial resolution and significantly enhanced penetration depth compared to conventional wide-field or confocal imaging techniques. These advantages have rendered it indispensable for structural and functional imaging in intact mammalian brains ^3,4^. However, conventional 2PM systems are typically limited to a FOV of up to ∼1 mm in diameter, which limits ability to simultaneously observe neuronal activity across multiple or widely distributed cortical areas. As systems neuroscience increasingly demands comprehensive, mesoscale interrogation of distributed neural circuits, the need for ultra-wide-field two-photon imaging, especially in deep layer, has become both evident and urgent.

Multiple technical approaches have been explored, with the most direct being the use of optical methods - particularly the iterative optimization of optical components such as objectives - to achieve the dual goals of high resolution and large field of view ^5–11^. However, these approaches have encountered a fundamental roadblock, with the key obstacle to achieving such systems lying in the fundamental trade-off dictated by the Smith-Helmholtz invariant, which states that the product of numerical aperture (NA) and FOV is conserved in optical systems. As a result, increasing the FOV necessarily leads to a reduction in achievable NA, thereby limiting spatial resolution and photon collection efficiency. This intrinsic trade-off poses a core challenge to the design of optical systems that aim to simultaneously achieve large FOVs and high-resolution, deep-tissue imaging. Therefore, current strategies have increasingly focused on incorporating complementary optical techniques using additional optical paths, micro-optical and mechanical components or optical modulation technologies to build upon existing systems, in order to circumvent the significant constraints imposed by conventional optical design. This, in turn, has increased the architectural complexity of two-photon microscopy systems designed for large FOV or extended-span imaging ^12–20^.

Efforts to expand FOV while preserving cellular resolution at depth are further hindered by chromatic aberration challenges inherent to multiphoton imaging. Through scattering tissue, most current wide-field multiphoton microscopes operate down to depths of ∼500 µm in the mouse brain, without employing techniques such as regenerative pulse amplification^21^, invasive methods to extend imaging depth ^17,19,22,23^, or the use of a more costly and technically complex three-photon excitation source^11,24,25^. Moreover, chromatic correction across a broad spectral range requires complex multi-element optics in the design of objectives, which introduce additional aberrations - coma, astigmatism, and field curvature - while increasing design complexity and the need for ultra-precise fabrication, inspection, assembly and adjustment of large-aperture components. As a result, these three aspects can rarely be minimized simultaneously. An integrative solution that combines ultra-wide FOV, high resolution, and deep-tissue imaging has been elusive.

Here, we present the ULTRA system (Ultra-wide, Layer-penetrating, Two-photon, Responsive Adaptive microscope; Fig. 1a, b), a next-generation imaging platform that overcomes these limitations through a synergistic excitation and collection strategy. First, we engineered a custom Ø8-mm FOV, 0.5 NA objective with 2 mm working distance optimized for multiple spectral ranges, and jointly optimized the objective-tube lens assembly for minimal aberration across the NIR excitation band, enabling efficient two-photon excitation across an 8-mm FOV (Fig. 1c left, d). A custom 2-axis adjustable head plate holder was designed to minimize the depth reduction caused by implantation surgery for the cranial window and headplate (Fig. 1c right). Second, we integrated a high-speed adaptive optics (AO) module^20,26^ employing a deformable mirror for real-time aberration correction, which substantially improves peripheral excitation at the edge of the FOV (Fig. 1e). Finally, we incorporated customized large-area photomultiplier tubes^13^ (PMTs, details see Fig. 1f) with enhanced current output range, significantly boosting the upper limit of signal dynamic range, which is particularly helpful for deep-tissue functional imaging^13^.

**Fig. 1.**
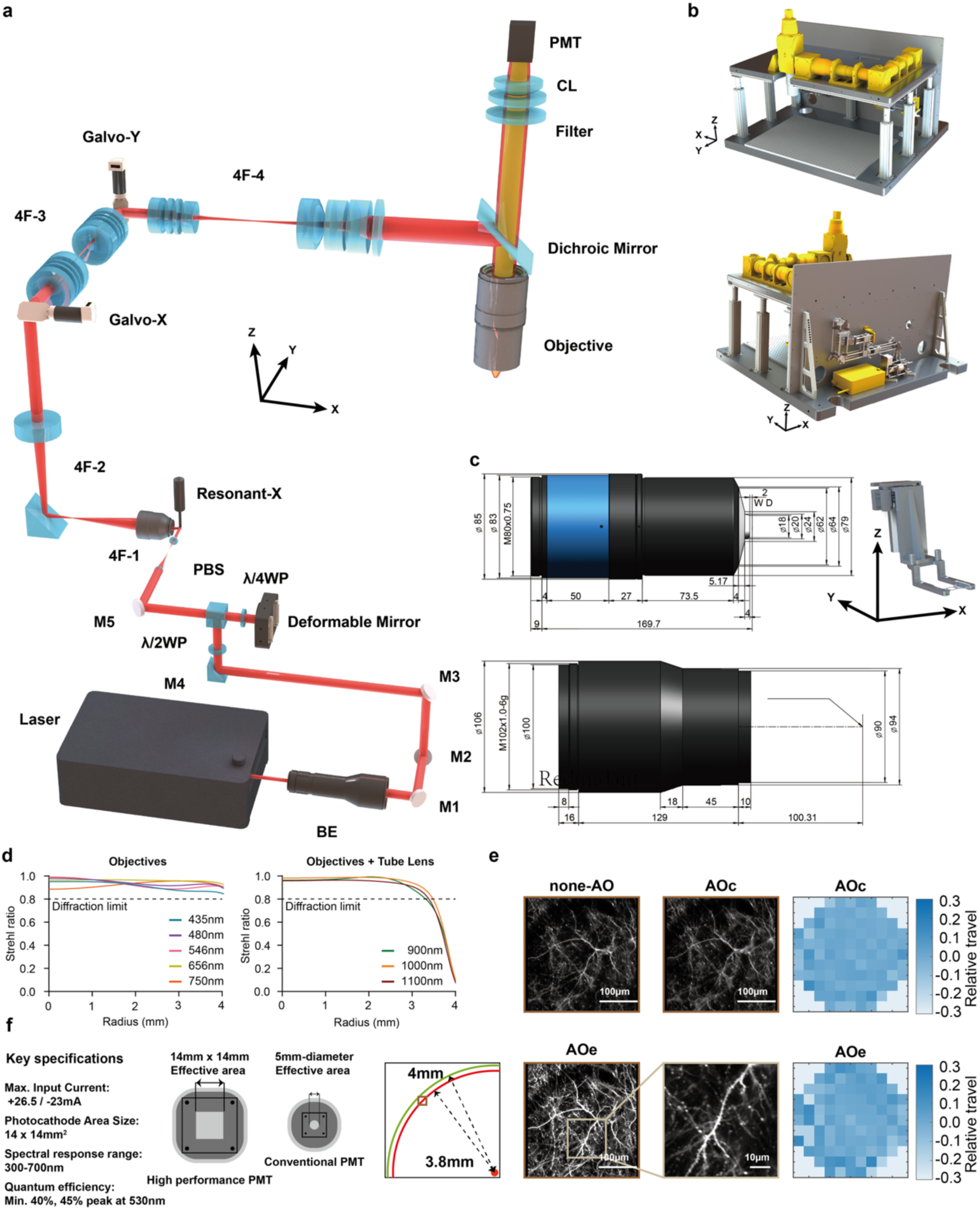
Schematic illustration of ULTRA highlighting key optical and mechanical components. **a**, Optical path layout of the ULTRA system. **b**, Mechanical structural design of ULTRA. **c**, Left: Custom-designed objectives and tube lens optimized specifically for the near-infrared (NIR) spectral range, ensuring exceptional photon collection efficiency and excitation performance. Right: Custom-built headplate holder with ±2.5° adjustable angles along both x and y axis, ensuring consistent imaging depth across multiple targeted cortical regions (unit: mm). **d**, Spectral optimization details for emission and excitation ranges. Left: Emission range optimization (435 nm, 480 nm, 546 nm, 656 nm, and 750 nm) ensures objectives maintain high Strehl ratios throughout the entire FOV radius (0–4 mm). Right: Excitation range optimization (900 nm, 1000 nm, and 1100 nm) with objectives and tube lens assembly. The integrated design of objectives and tube lens effectively corrects aberrations, although residual performance degradation at the FOV edge necessitates additional real-time correction via the deformable mirror when imaging across extensive spatial ranges. **e**, Validation of the adaptive deformable mirror’s performance for aberration correction at the periphery of the FOV. A dendritic spine located at a radial distance of 3.8 mm (maximum FOV radius: 4 mm, depth 83 μm, 50 mW) was imaged under three conditions: without adaptive optics correction (none-AO), adaptive optics correction optimized for the central region (AOc), and adaptive optics correction optimized for edge regions (AOe). Results demonstrate the capability to resolve individual dendritic spines at the edge of the FOV using appropriate adaptive optical correction. Corresponding phase diagrams illustrating the deformable mirror actuator configurations for AOc and AOe corrections are shown on the right. **f**, Key specifications and implementation of a high-performance photomultiplier tube (PMT) in ULTRA featuring an enlarged effective detection area (∼ 10 times) and enhanced output current compared to conventional PMTs (2 μA versus 50 μA). This design significantly improves visible fluorescence collection capability, extends the dynamic range for neural signal detection in functional imaging, and notably enhances the signal-to-noise ratio, particularly beneficial for deep-tissue imaging.

Together, these innovations enable high-resolution, high-SNR imaging across an Ø8-mm FOV and to depths approaching 900 µm in live brain tissue, with a spatial bandwidth product exceeding 5.27 × 10⁷. This platform thus provides a powerful and scalable solution for dissecting large-scale neural dynamics at single-cell resolution, bridging mesoscale systems neuroscience and deep-tissue cellular imaging.

## Results

### Optical design and layout of ULTRA

From a technical perspective, the design of optical objectives with a large FOV and high NA with 2 mm working distance inherently involves reconciling multiple conflicting requirements. Large FOVs typically exacerbate aberrations, however mesoscopic microscopy demands near diffraction-limited resolution, compounded by stringent apochromatic (APO) correction requirements. Balancing these three conflicting criteria presents significant challenges, particularly with regard to aberration management. As the FOV increases, field curvature and astigmatism at the edges become pronounced. Traditionally, these aberrations are mitigated by incorporating additional lens elements; however, this approach inherently introduces greater transmission losses and autofluorescence, negatively impacting the signal-to-noise ratio crucial in fluorescence imaging. Moreover, large FOV, cellular resolution, and deep tissue imaging together imply penetration of thicker biological samples, for which working distance is practically essential. Achieving a 2-mm working distance alongside a high NA with compact size is extremely difficult.

Our strategy for extending the practical limits included: (1) We defined the design envelope by precisely matching the optical etendue (G-value)^27^ with the physical dimensions of the intermediate image plane. This top-down approach ensured that the objective’s light-gathering capacity and spatial coverage were balanced to utilize the maximum theoretical capacity available. (2) To counteract the Petzval sum accumulation and to control field curvature inherent in high-etendue designs, we strategically redistributed optical power among lens groups. Specifically, we employed high-power negative lens elements positioned near aperture stops to effectively maintain high resolution across the entire ultra-wide FOV. (3) We utilized specialized high-cost dispersion glasses with precisely matched partial dispersion profiles. This allowed for the simultaneous suppression of primary and secondary spectra, achieving outstanding apochromatic (APO) performance across the target spectral range. (4) We moved beyond conventional standalone objective design by jointly optimizing the objective and tube lens as a single optical assembly. By treating the tube lens as an active compensator, we significantly increased the numbr of design variables (degrees of freedom) available. This synergistic approach allowed us to fine-tune residual aberrations, yielding near-diffraction-limited RMS wavefront performance across multiple excitation wavelengths and the entire imaging field (Fig. 1d and Supplementary Figs. 1-4). Consequently, this optimized design supports both ultra-wide FOV multiphoton imaging and multi-fluorescence imaging, with speckle illumination further enhancing optical sectioning (Supplementary Fig. 5). To do that, given the extensive spectral range requiring precise chromatic aberration correction, the objective incorporates cemented doublets and triplets composed of specialized optical glasses characterized by carefully selected relative dispersions. Glass selection is inherently constrained by the necessity to minimize intrinsic autofluorescence, essential for maintaining high fluorescence signal purity.

Typically, the front element of a high-numerical-aperture immersion objective is a hemispherical lens engineered as an aplanatic component conforming to the Abbe sine condition, thus minimizing axial spherical aberration and coma. Although reducing the radius of this hemispherical element is desirable to minimize spherical and chromatic aberrations (which scale proportionally with lens focal length), an inherent trade-off exists. Larger fields of view and flat-field imaging demand increased lens radii to reduce Petzval field curvature. To address this conflict, the hemispherical element in our flat-field apochromatic (APO) design is fabricated using high refractive-index, low-dispersion glass, effectively balancing the effects of increased radius. Moreover, the anterior surface of the hemispherical lens is intentionally formed into a negative meniscus curvature onto which a small plano-convex lens is cemented. This negative curvature substantially reduces the Petzval sum, while precise refractive index matching between the plano-convex lens and immersion medium mitigates reflection and refraction losses, significantly enhancing photon collection efficiency.

Subsequent low-dispersion lenses positioned behind the front hemispherical element further decrease the convergence angle of incoming rays, facilitating more effective correction of axial and lateral chromatic aberrations and field curvature in subsequent lens groups. Residual optical aberrations, including spherical aberration, are further minimized through strategic employment of low-dispersion positive lenses and high-dispersion negative lenses, combined with thicker lens elements and meticulous optimization of lens curvatures. Near the exit pupil, a strategically placed concave lens reduces ray height and introduces additional negative curvature, substantially diminishing the Petzval sum and ensuring superior flat-field performance across the entire imaging field.

The layout of ULTRA is set out below (Fig. 1a, b). A 920 nm fiber-based femtosecond laser (ALCOR 920-4-XSight, Spark lasers) serves as the source, and its beam is expanded to approximately 14 mm using a beam expander (BE, FBE-10X-B, LBTEK). The expanded beam is then reflected by a half-wave plate (λ/2WP, WPA2420-650-1100, Union Optic) and a polarizing beam splitter (PBS, MPBS643, LBTEK) to ensure vertical entry into a deformable mirror (DM97-15, ALPAO). After vertical reflection from the DM, the beam passes through a quarter-wave plate (λ/4WP, AQWP10M-980, Thorlabs) to adjust its polarization, allowing it to pass straight through the PBS. The DM is optically conjugated with a resonant scanning mirror (Resonant-X, 8 kHz CRS, Cambridge Technology) via a 4F-1 system (AC254-200-B and AC254-075-B, Thorlabs). This resonant scanner is then optically conjugated with the galvanometer mirror Galvo-X (QS20X-AG, Thorlabs) through another 4F-2 system (LSM54-1050 and TTL200MP, Thorlabs). Galvo-X and its counterpart Galvo-Y (QS20Y-AG, Thorlabs) are optically conjugated using a custom-designed 4F-3 system comprised of two lens groups (each with f = 90 mm). Finally, Gavlo-Y is optically conjugated through an additional custom-designed 4F-4 system—consisting of a 90 mm lens group and a 200 mm lens group—with the entrance pupil of a custom-designed large field-of-view objective (object field diameter: 8 mm, NA: 0.5, working distance: 2 mm). A custom-designed dichroic mirror is used to separate the near-infrared femtosecond laser light from the visible fluorescence. The collection lens (CL) is a custom-designed large-aperture lens assembly (F = 60 mm), and an infrared cut-off filter (ET700SP-2P8, OD = 8, two-inch) is placed before the PMTs to further block any residual infrared laser light. To accommodate large-field fluorescence detection, a GaAsP PMT with a 14 mm x 14mm detection aperture (H15460-40, Hamamatsu) is employed.

### Performance characterization of ULTRA

To characterize the spatial resolution of ULTRA, we performed z-stack acquisitions of 0.5 μm fluorescent microspheres distributed at multiple positions within an 8 mm-diameter FOV. The FOV was segmented into 1 center spot and 3 rings based on the radial distance from the center (Fig. 2a). Each ring was further subdivided into eight measurement positions, spaced at 45° intervals, to comprehensively assess spatial resolution across the entire field.

**Fig. 2.**
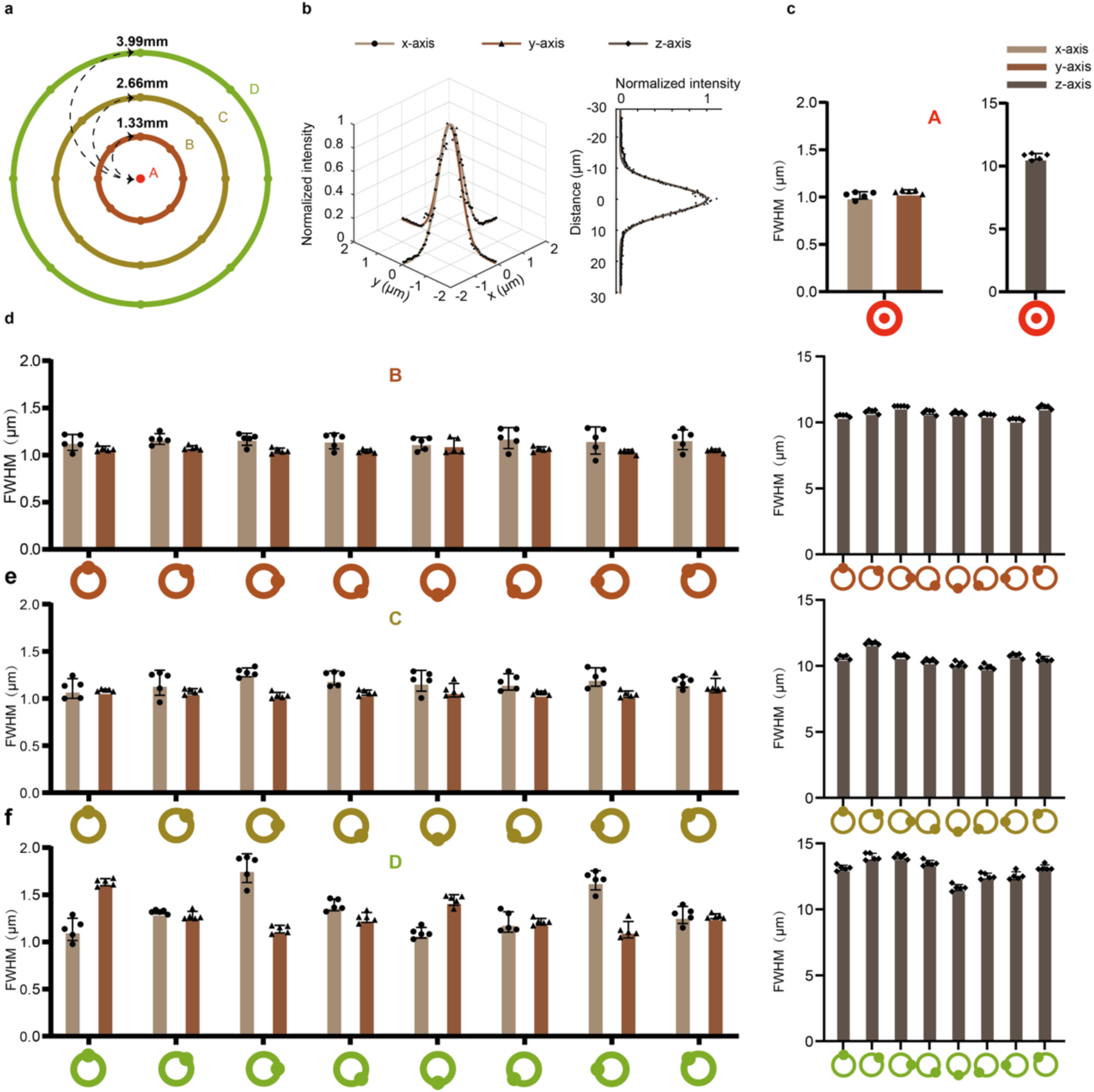
Spatial resolution characterization of ULTRA. **a**, Schematic diagram of spatial resolution measurement locations across the FOV. **b**, Example of measured data and corresponding Gaussian fitting results from center point A, imaged with voxel size x: ∼0.06 μm y: ∼0.06 μm z: ∼0.27 μm. **c**, Summary of lateral and axial resolution characterization at center point A (n=5). **d**, **e**, **f**, Summary of lateral and axial resolution measurements taken at points distributed at 45° intervals along three concentric rings (radii: 1.33 mm, 2.66 mm, and 3.99 mm from the center, n=5 each).

At the central point A (Fig. 2b, c), lateral resolution was determined to be 1.05 ± 0.05 μm in the x-axis and 1.05 ± 0.02 μm in the y-axis, with axial (z-axis) resolution of 10.88 ± 0.21 μm (mean ± SD). For the B, C and D ring (Fig. 2d), the measured resolutions were as follows (Fig. 2e, f): Ring B: X = 1.15 ± 0.09 μm, Y = 1.06 ± 0.04 μm, Z = 10.77 ± 0.32 μm, Ring C: X = 1.19 ± 0.10 μm, Y = 1.07 ± 0.05 μm, Z = 10.62 ± 0.54 μm, Ring D: X = 1.36 ± 0.25 μm, Y = 1.30 ± 0.17 μm, Z = 13.06 ± 0.79 μm. (mean ± sd, see Supplementary Table. 1 for details). Furthermore, we performed additional measurements at depths of up to 1000 μm (Supplementary Fig. 6). Our analysis shows that the system maintains a lateral resolution of 1.19 ± 0.06 μm (mean ± SD) for the x-axis and 1.20 ± 0.05 μm for the y-axis, with axial resolution of 11.70 ± 0.37 μm, within the depth range of 900-1000 μm, confirming that it remains sufficient for resolving cellular structures deep within tissue. These measurements provide a comprehensive characterization of the spatial resolution across the imaging field, demonstrating ULTRA’s competence at resolving cellular structure throughout a 50.27 mm^2^ FOV.

### Flexible in vivo two-photon imaging of neuronal morphology and cerebral vasculature

To comprehensively demonstrate the in vivo imaging performance of our system, we conducted a series of experiments. First, we surgically implanted a custom titanium headplate and an 8-mm diameter cranial window into the skulls of transgenic mice expressing GAD67-GFP in GABAergic interneurons under control of the GAD67 promoter. The head was fixed via the cranial head plate to an adjustable holder that permitted ±2.5° adjustments in two orthogonal directions, compensating for any horizontal misalignment introduced during the sequential implantation procedures between headplate and cranial window, ensuring the targeted cortical areas remained perpendicular to the surface of objective lens, and maintaining the depth across different regions. Using the 8-mm cranial window, we acquired images of the entire underlying tissue (Fig. 3a, left; pixel dimensions 4400×4800, frame rate 0.83 Hz). Ten distinct regions were selected, and upon magnification, the somata of GABAergic neurons were clearly resolved throughout the entire FOV.

**Fig. 3.**
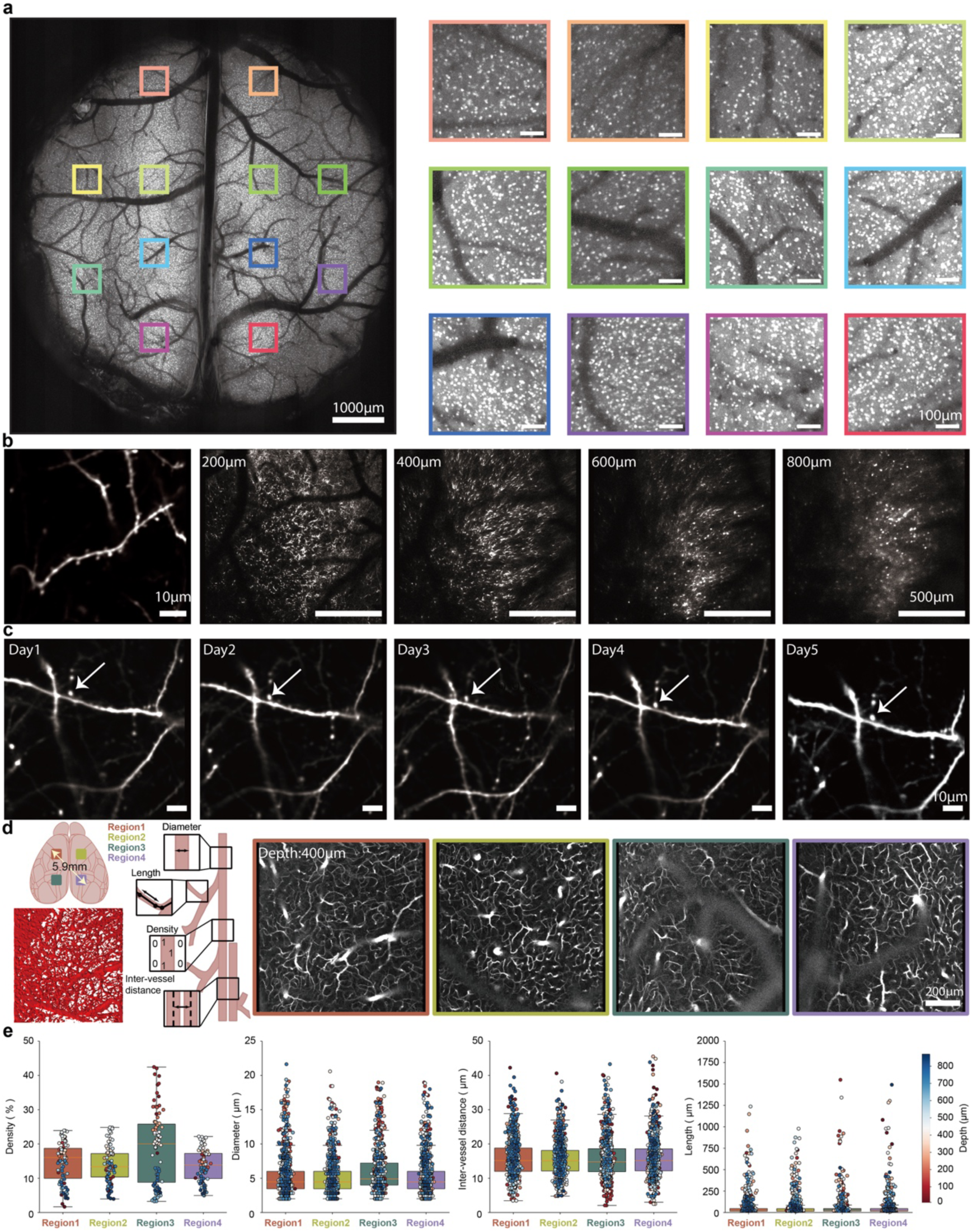
Multi-regime ULTRA imaging of a broad range of cortical structures. **a**, Ultra-wide-field two-photon imaging of GFP-labeled GABAergic interneuron somata performed through an 8-mm cranial window. The image represents a maximum-intensity projection of a 100-μm-thick subcortical z-stack (total z-stack volume: 300 μm). The right panel shows 10 digital zoomed-in sub-fields distributed across the entire FOV. **b**, Left: Dendritic spines along an individual dendrite in the cortex of a Thy1-GFP transgenic mouse are resolved at a depth of 145 μm using 40 mW laser power. Right: Dendritic and somatic structures are delineated at various depths within a volume of 1.65 mm × 1.65 mm x 0.9 mm, power 107.5 mW. **c**, The expansive FOV enables longitudinal registration and imaging of the same dendritic spine, as demonstrated by continuous observation over 5 days at a depth of 70 μm. **d**, Left: schematic diagram of the simultaneous imaging of vasculature across 4 distinct cortical regions in the mouse, along with an example of vascular reconstruction in Region 1 (top view) and the diagram of 4 vascular measurement parameters—vessel diameter, length, density, and inter-vessel distance. Right: Vascular imaging from the four regions (shown at depth of 400 μm, reaching up to 870 μm) was performed following tail-vein injection to label the vasculature. The maximum inter-regional span was 5.9 mm, the imaging frame rate was 6.02 Hz, and each FOV measured 1 mm × 1 mm. **e**, Boxplots summarize the statistical analysis of vascular parameters measured across the 4 regions.

Next, to assess the system’s ability to resolve fine neuronal structures in vivo, we imaged dendrites and dendritic spines in transgenic mice expressing Thy1-GFP under the control of the Thy1 promoter (Fig. 3b). Initially, individual dendritic spines were imaged (Fig. 3b, left), revealing clear delineation of single spines. Subsequently, z-stack imaging was performed over a volume of 1.65×1.65×0.9 mm³ (Fig. 3b, right, see whole z-stack imaging in Supplementary Fig. 27). To evaluate long-term stability in imaging fine structures such as dendrites and dendritic spines, we repeatedly imaged the same targets over 5 days (Fig. 3c). These results not only confirm the high optical resolution of our system but also its robust capability for chronic, longitudinal deep tissue imaging.

To evaluate imaging performance over a large field of view in deep tissue, we conducted simultaneous multi-area in vivo imaging of mouse cerebral vasculature labeled with i.v. injection of Dextran-Texas Red, at a frame rate of 6.02 Hz (Fig. 3d). Z-stacks of images were acquired across four regions spanning both hemispheres, with the maximum distance between these 4 independent regions being over 5.9 mm, each with an imaging volume of 1×1×0.87 mm³. This allowed us to perform a quantitative analysis of cortical vascular architecture in wild-type mice (Fig. 3e, Supplementary Video 1). Collectively, these results demonstrate that the system achieves excellent spatial resolution across both superficial and deep tissues, even under large-field-of-view and wide-span imaging conditions.

### Flexible in vivo two-photon calcium imaging delivering across diverse spatial scales and depths

Following structural imaging of cortical tissues in mice, we performed a series of in vivo calcium imaging experiments in awake animals. In these experiments, transgenic mice expressing the genetically encoded calcium indicator GCaMP6s in the control of Thy1 promoter in cortical neurons were utilized. Initially, we replicated conventional state of the art wide-field two-photon imaging, which typically employs a FOV of 3-5 mm diameter ^10,12,13^ (Fig. 4a). Therefore, a 5-mm-diameter cranial window was implanted onto the mouse skull, enabling simultaneous observation of neuronal activity in layers 2/3 of both hemispheres within a 4 × 4 mm² FOV (Fig. 4a, left, Supplementary Video 2). At an imaging depth of approximately 190 μm and a pixel ratio of 0.35 pixels/μm, a total of 10,026 neurons were detected throughout the entire FOV. Representative ΔF/F calcium traces with prominent signal transients from both hemispheres are shown (Fig. 4a, right). Analysis of the raw data yielded a signal-to-noise ratio (SNR) of 5.09 ± 1.05 (mean ± s.d). Notably, the calcium signals acquired by this system exhibit exceptionally high Δf/f values, substantially improving the sensitivity and quality of calcium signal detection.

**Fig. 4.**
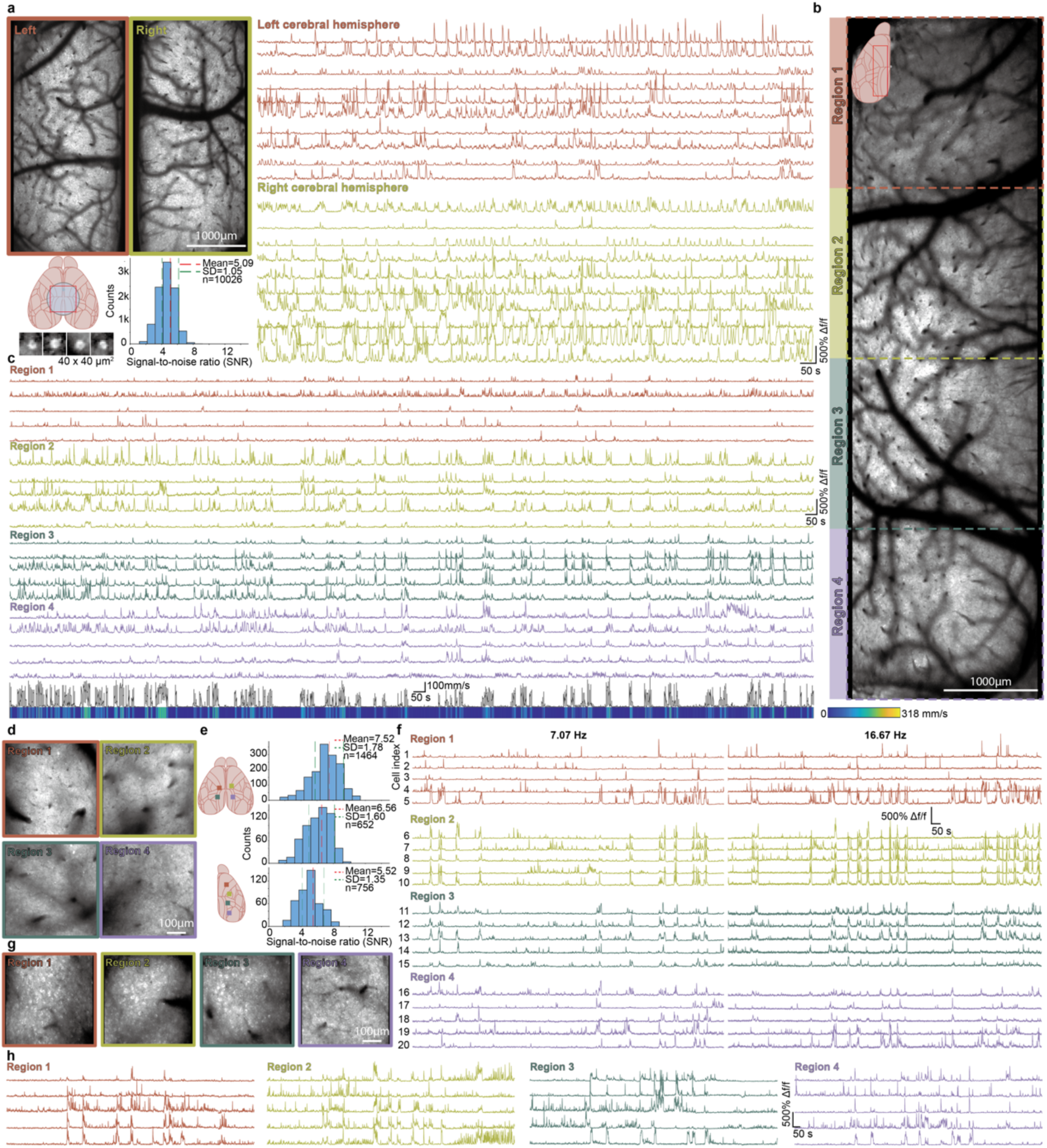
ULTRA calcium imaging of neuronal activity of locomotory mice at multiple depths spanning an ultra-wide FOV. **a**, Conventional wide-field two-photon calcium imaging (Thy1-GCaMP6s). Neurons displayed (lower left) are cropped from the full FOV. A schematic illustrates cranial window placement (lower left), alongside representative calcium signals from both hemispheres and corresponding SNR measurements (mean ± sd). Imaging was performed at a depth of 240 μm with pixel size of 1400 × 1400 pixels over 4 x 4 mm^2^ and frame rate of 5.46 Hz. **b**, High-power, long-duration imaging in the cortex of locomotory mice across an extensive continuous FOV spanning 7.28 mm across 10 cortical regions. Imaging was conducted at a depth of 259 μm using 300 mW laser power over 1 hour, at frame rate of 6.36 Hz with pixel dimensions of 700 × 2450 pixels. **c**, Calcium signal traces extracted from four discrete regions within the FOV in **b** are shown, with the corresponding locomotion speed trace displayed beneath. **d**, Shallow-layer two-photon calcium imaging across 4 independent regions under varying acquisition rates. Each region was imaged at depth of 250 μm within 500 x 500 μm^2^ FOV (imaging sites are indicated in the upper left of **e**). Two acquisition conditions were employed: 7.07 Hz (500 × 500 pixels; SNR = 7.52 ± 1.78, statistics shown in the upper right of **e**) and 16.67 Hz (170 × 170 pixels; SNR = 6.56 ± 1.60, statistics shown in the middle right of panel **e**). **e**, Diagram of Imaging sites in experiments of **d** and **g**, and the corresponding SNR measurements **f**, Calcium signal traces from identical neurons acquired at two imaging frame rates are presented; traces on the left were captured at 7.07 Hz, and those on the right at 16.67 Hz. **g**, Deep-layer two-photon calcium imaging across four independent regions was performed at a depth of 519 μm using 250 mW laser power. The imaging sites, as shown in the lower left of panel **e**, span a maximum distance of 5.56 mm; corresponding calcium signal traces from these regions are presented in **h** (7.07 frame rate, SNR = 5.52 ± 1.35, statistics shown in the lower right of **e**).

To demonstrate the system’s capability for monitoring neuronal population dynamics over extended spatial scales, we next performed calcium imaging over a larger FOV. Utilizing a customized curved cranial window^28,29^ (∼7 × 8 mm²), we imaged neuronal activity in layers 2/3 within a 2 × 7 mm² region in the left hemisphere (Fig. 4b, Supplementary Video 3). Continuous recordings were acquired over one hour, during which laser power was intentionally elevated (300 mW) to assess potential photodamage under high-intensity illumination. At an imaging depth of 258 μm, high SNR calcium signals were extracted across the entire FOV (Fig. 4c), while concurrent measurements of the mouse’s locomotor velocity were recorded (Fig. 4b, bottom). Notably, no photobleaching, laser-induced tissue damage, or neuronal and behavioral abnormalities were observed throughout the session.

After validating very-wide-FOV calcium imaging, we performed simultaneous calcium recordings from four distinct regions spanning both hemispheres under a 5 mm cranial window at acquisition rates of 7.07 Hz and 16.67 Hz respectively (Fig. 4d, Supplementary Video 4 and, Supplementary Video 5). Calcium traces with different frame rate samplings applied to the same neurons, extracted from each region, are shown (Fig. 4f) with SNR of 7.52 ± 1.78 with 7.07 Hz and 6.56 ± 1.60 with 16,67 Hz respectively.

Finally, deep tissue calcium imaging was conducted in four regions within a single hemisphere at an imaging depth of 519 μm (Fig. 4g, h, Supplementary Video 6), achieving an SNR of 5.52 ± 1.35. These experiments demonstrate that our system maintains exceptional spatiotemporal resolution, enabling high-fidelity and versatile calcium imaging of superficial cortical neurons under diverse FOV configurations as well as capturing deep-layer neuronal activity over extended spatial scales.

### Flexible in vivo two-photon calcium imaging characterizes the neuronal coactivity network for mouse locomotion across multiple scales

Having demonstrated the high-quality cross-scale cortical calcium imaging capability of our system in moving mice, we further examined its flexibility in detecting cortical calcium signals during mouse locomotion by conducting three distinct experiments targeting different neuronal networks^30^. Real-time locomotion data were captured using an air-floating head-fixed mouse behaviour platform^31,32^ (Fig. 5a, b). In the first experiment, the animal underwent functional neuronal imaging across an expansive field of view (FOV) of 2 mm × 7 mm (spanning up to 7.28 mm), covering a total of 7,931 neurons across 10 brain regions in the left hemisphere, including motor, somatosensory, visual, and retrosplenial cortex. Comparing the functional neuronal network connectivity between stationary and moving states within the left cortical hemisphere revealed a marked increase in the number of intra-regional neuronal connections (from 82 to 684) and inter-regional connections (from 143 to 689) at a correlation threshold > 0.3 (Fig. 5c, d). These results clearly demonstrate that locomotion activates significantly more neurons, expands the spatial extent of neuronal network coupling to a more global scale, and enhances functional connectivity strength (Fig. 5d, e).

**Fig. 5.**
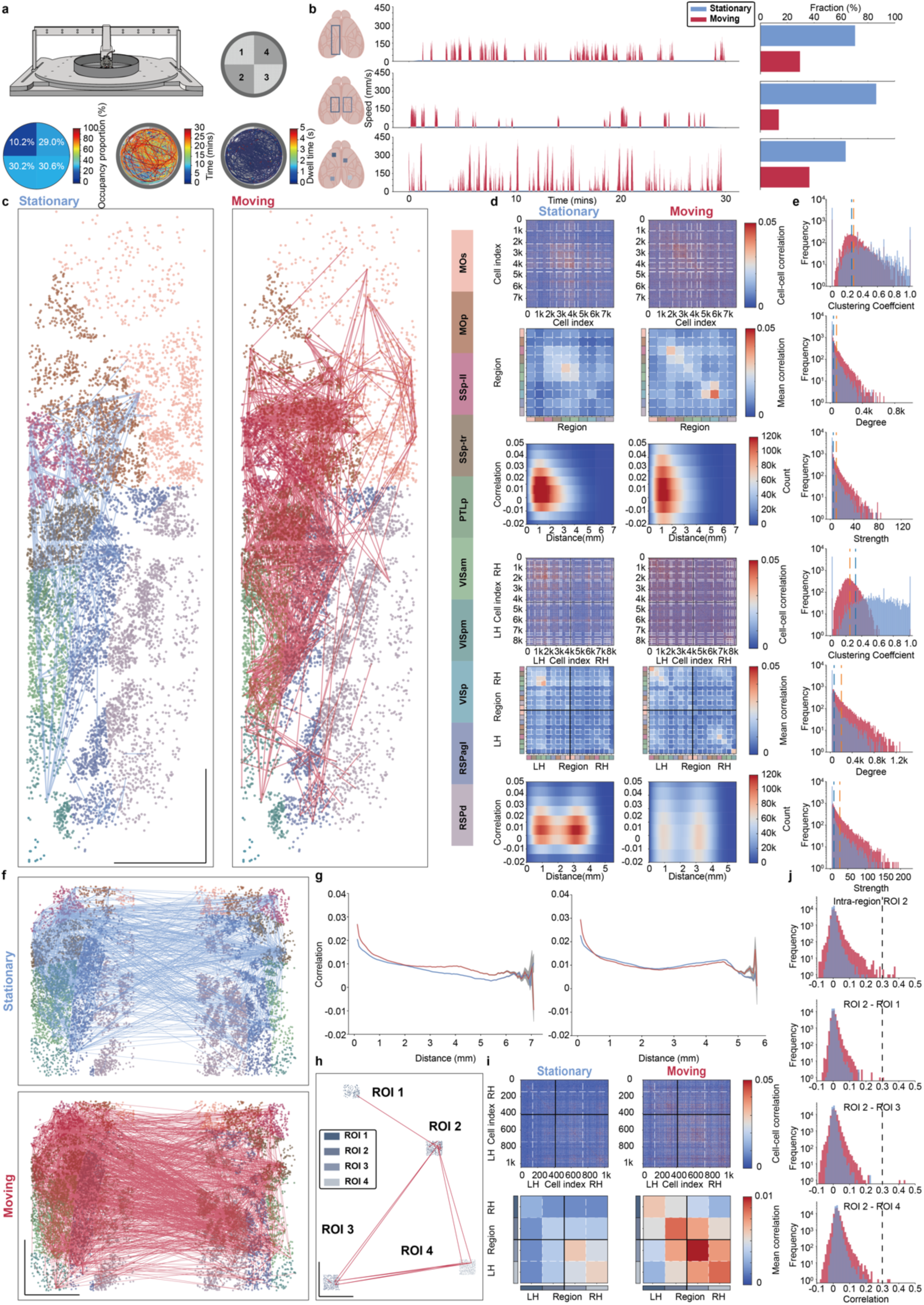
Functional connectivity assessments on neuronal ensembles over regions in locomotory mice via ULTRA imaging. **a**, The schematic diagram of air-floating platform for animal locomotion monitoring and examples of locomotion recording from the air-floating platform in 30min, top right, the carbon fiber open-field chamber with 4 areas. Bottom left, occupancy proportion in each area. Bottom middle, animal locomotion trajectory and bottom right, dwell time along the locomotion trajectory. **b**, Imaging FOV demonstrations and speed measurements of animals in the following experiments, from top to bottom, the first experiment to the third, and corresponding locomotion speed traces and fractions in different states (stationary (blue) and moving (red), experiment # 1, 70.49% with mean ± sd = 1.2 ± 1.26 mm/s for stationary and 29.51% with mean ± sd = 70.72 ± 44.99 mm/s for moving, experiment # 2, 86.22% with mean ± sd = 0.98 ± 0.96 mm/s for stationary and 13.78% with mean ± sd = 79 ± 46.44 mm/s for moving, experiment # 3, 63.48% with mean ± sd = 1.59 ± 1.68 mm/s for stationary and 36.52% with mean= 89.60 ± 89.13 mm/s for moving). **c**, First experiment (2 mm x 7 mm FOV, 6.36 Hz at 250 μm depth (Thy1-GCaMP6s)), changes in functional connectivity among 10 cortical regions on left hemisphere across behavioral periods (connections shown with the same threshold in both states (0.3, Pearson Correlation Coefficient) in all experiments, blue: stationary, red: moving), right, brain regions according to Allen Mouse Common Coordinate Framework with corresponding color code shown in the left (MOs: secondary motor area, Mop: primary motor area, SSp-ii: primary somatosensory area, lower limb, SSp-tr: primary somatosensory area, trunk, PTLp: posterior parietal association area (VIsa Anterior area), VISam: anteromedial visual area, VISpm: posteromedial visual area, VISP: primary visual area, RSPagl: retrosplenial area, lateral agranular part, RSPd: retrosplenial area, dorsal part). **d**, 7913 cells among 10 regions extracted from experiment # 1 were used to measure over 31.30 million cross-correlations pairs, plotted by cells and regions and the distribution of correlation as a function of distance between neurons ranging from 0 μm to 7048 μm (top 3, LH: Left Hemisphere, RH: Right Hemisphere, white dashed lines: brain region boundaries), and same analysis for experiment # 2 (shown in **f,** bottom 3, 8601 cells, 18 regions bilaterally, over 36.98 million cross-correlations pairs, neurons ranging from 0 μm to 5.551 μm, black lines: hemisphere boundaries). **e**, Neuronal networks were assessed regarding correlations measured with graph theory (threshold=0.1 in clustering coefficient, degree and strength calculations) in experiment # 1 (top 3) and # 2 (bottom 3) with mean value presented in dark blue (stationary) and orange (moving) dashed lines. **f**, Second experiment (2 x 2 mm x 3.5 mm FOV, 6.02 Hz at 225 μm depth (Thy1-GCaMP6s)), same in **c**, 18 cortical regions on bilateral cerebral hemispheres. **g**, Correlations in **d** plotted as a function of distance between neurons, experiment # 1 (left), experiment # 2 (right). **h**, Third experiment (4 x 0.5 mm x 0.5 mm FOV, 16.67 Hz at 598 μm depth (Thy1-GCaMP6f)), same in **c** and **f**, 4-region simultaneous calcium imaging in deep layer with higher frame rate with maximum connection distance of 6687 μm. **i**, Same measurements in **d** (top 2). **j**, Examples of distribution of correlation intra-region of Region 2 and between Region 2 and the other regions, and functional connectivity beyond dashed line (threshold 0.3) were shown in **h**. Scale bar: 1mm.

We subsequently expanded the imaging FOV to cover bilateral cortical hemispheres, acquiring 30-minute recordings spanning a maximum 6.10 mm, each with an FOV of 2 mm × 3.5 mm. Visualizations (Fig. 5f) and quantitative analyses (Fig. 5d, e) of neuronal network connectivity confirmed that networks involving 18 cortical regions from both hemispheres exhibited similarly enhanced connectivity during locomotion consistent with previous experimental results. This enhancement was evident in increased intra-regional connections (from 244 to 853), inter-regional connections (from 689 to 2,015), notably pronounced interhemispheric connectivity, spatial distribution, and connection strength (Fig. 5d, e, f). To elucidate how neuronal correlation varies with distance, we also measured Pearson correlation coefficients between neuronal time series as a function of inter-neuronal distance, revealing relatively high correlations within 0–300 μm, which gradually decreased at greater distances (Fig. 5g).

These two experiments demonstrate the capability to monitor large-scale superficial (layer 2/3) cortical neuronal networks. To further validate the performance of our system, we conducted rapid acquisition imaging of deep cortical neuronal activity using a faster calcium indicator (Thy1-GCaMP6f transgenic mice). We imaged four discrete cortical areas distributed across both hemispheres, spanning over 6.6 mm with imaging depths up to ∼ 600 μm. Analysis of the collected data from 1,044 neurons (Fig. 5h–j, Supplementary Fig. 8) revealed consistent enhancement in neuronal network connectivity and connection strengthnduring locomotion (Supplementary Fig. 9). Taken together, these results, achieved through ultra-wide FOV imaging with high-efficiency deep tissue penetration and exceptional signal-to-noise ratio, underscore the flexibility and superior optical performance of the ULTRA system, facilitating a series of experimental measurements across both superficial and deep cortical layers.

## Methods

### Mice

All mice (8-16 weeks) used in this study were bred in the animal facility of Suzhou Institute of Biomedical Engineering and Technology and approved in accordance with the Institutional Animal Care and Use Committee guidelines at Suzhou Institute of Biomedical Engineering. Mice were housed in individually ventilated cages under a 12-hr light/12-hr dark cycle with normal food and water. C57BL/6J mice were used for vascular imaging, GAD67-GFP knock-in mice ^33,34^ and Thy1-EGFP mice (Jackson Labs stock # 007788)^35^ were used for GABAergic interneuron imaging and subcellular structure imaging. For functional imaging, both Thy1-GCaMP6s C57BL/6J-Tg(Thy1-GCaMP6s)GP4.3Dkim/J and Thy1-GCaMPf C57BL/6J-Tg(Thy1-GCaMP6f)GP5.17Dkim/J (Jackson Labs stock # 025393 and # 024275)^36^ were used to perform calcium signal recording.

### Surgery

Mice were anaesthetised with 1.5-2% isoflurane with body temperature maintained at 37 ℃. Analgesia was administered pre-operatively with Carprofen (5 mg/kg). Anaesthetic depth was assessed every 10 min for the duration of the experiment. Cover the eyes with ophthalmic ointment and depilated and disinfect the surgical area. Vetbond tissue adhesive was used to seal the skin wound around the skull. After removing the mouse’s scalp, the membrane was removed, the skull surface was cleaned. Craniotomy was made manually with 0.5 mm and drill bit, and then the loose skull segment removed. Dental adhesive resin cement was used to glue the cranial window and headplate and ensure all surgical areas were covered to avoid direct exposure. Mice were given carprofen (5 mg/kg/24 hrs) in oral water for 3 days after surgeries. Mice were allowed 1 week to recover before imaging began.

### In vivo two-photon imaging

All experiments were conducted on the customized ULTRA system. Both the microscope and image acquisition were controlled by custom-written software using LabVIEW 2020 (National Instruments). For two-photon imaging of morphology, mice were anaesthetised with 1.5-2% isoflurane with body temperature maintained at 37 ℃. Anaesthetic depth was then assessed every 10 min for the duration of the experiment. Eyes were covered with ophthalmic ointment prior to imaging.

For two-photon imaging of calcium signals, animals were habituated for 2 sessions on the day before imaging (10 minutes per session, 4 hr intervals between sessions). Animals were then placed in the air-floating chamber for 5 minutes before imaging started. The custom-built headplate holder was used to reduce any angular deviation caused by the implantations of the cranial window and headplate during surgery, which can induce a difference in accessible tissue depth between areas when imaging over a large span. Imaging was normally performed with less than 250 mW emitted from the front of the objective. Imaging parameters: pixel ratio = 0.35, unit pixels/μm for wide field and multi-region imaging, with a frame rate of ∼17 Hz, and pixel ratio = 1 when conducting multi-region imaging, with a frame rate of ∼7 Hz.

### Data analysis for neuronal calcium imaging

Calcium movies were segmented and analyzed using custom scripts in MATLAB (Mathworks) and Python. Cellular ROIs and corresponding fluorescence time series were extracted from imaging stacks using Suite2p^37^ (https://suite2p.readthedocs.io/). The neuropil component was subtracted from the neuronal signals. Relative fluorescence changes (Δf/f) were used in our analysis, with Δf/f = (f − f0)/f0 calculated^38,39^, where the fluorescence changes (f) average the pixel values in each specified ROI, and baseline fluorescence f0 is estimated as the 15th percentile of the entire fluorescence recording. Pearson correlation of calcium traces between neurons was calculated using Python, and graph theoretic analysis ^40–43^ was applied to the data shown in Fig. 5 using the NetworkX package in Python.

### Vascular imaging analysis

Vascular imaging stacks were segmented using the *DeepVess* pipeline^44^ and quantified using MATLAB and Python scripts. Images consisted of 600 x 600 pixels (1000 μm x 1000 μm) with a z-axis step size of 10 μm. As a preprocessing step, images were intensity-matched using adaptive histogram equalization and subjected to crosstalk removal, dura removal, normalization, noise reduction, and motion correction. Vessel segmentation was then performed using the DeepVess 3D Convolutional Neural Network (CNN) model, which consists of six 3D convolutional layers with ReLU activation and max-pooling, followed by a softmax layer for classification. After segmentation, the diameter, length, inter-vessel distance and density were computed^45–48^.

The diameter *D* of a vessel segment was determined by first computing the radius (*r_j_*) at each point along the vessel centerline as the shortest distance from that point (*v_j_*) to the nearest background voxel:

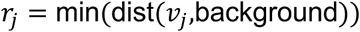

Once all *r_j_* values are obtained, their median is calculated. The diameter is then given by twice this median radius:

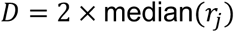

The length *L* of vessel segments was computed by summing the Euclidean distances between consecutive points along the vessel centerline:

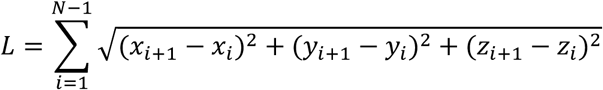

The inter-vessel distance (*IVD*) was determined as the minimum Euclidean distance between the midpoints of two closest vessels:

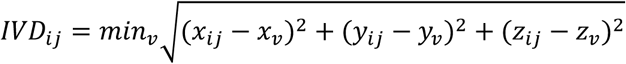

The vascular density ρ was defined as the ratio of the total volume occupied by the vessels *V_v_* to the total volume of the region of interest *V_RoI_*:

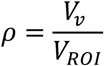

### Adaptive optics

System aberrations were corrected using AO with mixed pollen grain samples (Mixed Pollen Grains w.m., Carolina). Given the nonlinear nature of two-photon excitation, fluorescence intensity scales with the square of the spatial photon density; consequently, under constant average laser power, a minimized focal volume yields higher photon density and enhanced fluorescence excitation. We therefore implemented a sensorless AO approach, utilizing the mean fluorescence intensity of the pollen images as the feedback metric to optimize for minimal focal spot size (indicative of minimized aberrations). The coefficients of the first 200 Zernike modes were optimized sequentially. For each mode, the coefficient was modulated continuously within a preset range (21 steps per cycle) and repeated over five cycles to mitigate error. The coefficient corresponding to the peak fluorescence intensity was selected as the optimal compensation value and incorporated into subsequent iterations. With each coefficient transition corresponding to a single image frame, the high switching speed of the deformable mirror combined with a high-frame-rate imaging mode (133 Hz for local ROI scanning) significantly accelerated the calibration. The entire optimization process, comprising approximately 20,000 coefficient transitions, was completed in approximately 3 minutes.

### Electronic control system and software description

The central control system was built on a PXI platform (PXIe-1082, National Instruments) utilizing an FPGA multifunction card (PXI-7851R, National Instruments) to generate synchronized acquisition timing and Galvanometer scanning waveforms. Fluorescence signals were acquired using a high-speed digital oscilloscope acquisition card (PXIe-5122, National Instruments; 100 MHz sampling rate), while data communication between the PXI chassis and the host server (Precision T7920, Dell) was enabled via a PXI-to-PCI communication card (PXIe-PCIe 8398, National Instruments). The imaging software, developed on the LabVIEW platform (LabVIEW 2020), encompasses functionalities for electronic translation stage control, adaptive optical correction, scan mode configuration, image acquisition, reconstruction, display, and storage. Scanning modalities include a single-region high-speed mode (typically employed for adaptive optics correction), full-field stripe scanning, and ROI scanning. In vivo image processing further integrates resonant scan sinusoidal distortion correction and real-time sliding average display.

### Resolution measurement

Optical resolution was quantified using 0.5 μm fluorescent microspheres (FSDG003, Bangs Labs) at depth of ∼ 30 μm in a 0.75% agarose gel. Prior to measurement, pixel dimensions were calibrated with a Ronchi ruling test target (R1L3S12N, Thorlabs), yielding pixel sizes of 0.0564 μm in the X-direction and 0.0557 μm in the Y-direction. Axial resolution was assessed by modulating the microsphere sample’s position with a motorized stage in 0.27 μm increments. A custom MATLAB script was employed to fit the acquired data with a Gaussian function, with the full width at half maximum (FWHM) of the fitted curve serving as the resolution metric.

### Key Component Specifications

A femtosecond laser (ALCOR 920-4-XSight, Spark Lasers) was employed as the excitation source with a central wavelength of 920 nm, pulse duration of 119 fs, and an average power of ∼4 W. These parameters were optimized to efficiently induce non-linear two-photon absorption while minimizing scattering losses in deep tissue. To actively correct wavefront distortions caused by refractive index inhomogeneities across the large field of view (FOV), a deformable mirror (DM97-15, ALPAO) was incorporated into the system. It features a 13.5 mm clear aperture and 97 micro-mirrors capable of modulating the optical phase with a settling time of 0.8 ms. For beam scanning, a resonant scanner (8 kHz CRS, Cambridge Technology) with an 8 kHz resonance frequency was utilized for high-speed fast-axis scanning. This was paired with a galvanometer scanner (QS20XY-AG, Thorlabs) possessing a 20 mm clear aperture. This large aperture is physically essential to minimize vignetting and fully fill the back pupil of the high-NA objective, thereby preserving the system’s optical resolution. A custom-designed, high-resolution wide-field microscope objective was implemented, offering an 8 mm object-side FOV and a numerical aperture (NA) of 0.5. The objective utilizes a water-immersion configuration to match the refractive index of biological tissue (∼1.33), significantly reducing spherical aberrations at the interface. To maximize the detection of ballistic and multiply-scattered photons via the photoelectric effect, a photomultiplier tube (H15460-40, Hamamatsu) was used. It is equipped with a GaAsP photocathode exhibiting a high quantum efficiency of 45% at 520 nm and a large photosensitive area of 14 mm × 14 mm, which increases the collection solid angle for diffuse signals. A three-dimensional motorized translation stage (XWJ-200R-2G and EVSN30P, Sanying MotionControl Instruments Ltd) provided high-precision sample positioning. Finally, the system was integrated with an air-floating mobile cage system (Neurotar tracker), which utilizes aerostatic lubrication to eliminate friction, allowing the head-fixed animal to navigate naturally during imaging.

## Discussion

We have here demonstrated ULTRA, a very wide field of view multiphoton microscopy system that overcomes the limited deep-tissue imaging performance with NA 0.5 optical systems. This system achieves a remarkable lateral imaging diameter of 8 mm while maintaining effective imaging depths approaching nearly 1 mm in brain tissue, performance that, in certain aspects, catches up with or even surpasses traditional high-NA two-photon microscopes. The exceptional capability of this system arises from an innovative optical design integrated with adaptive optics and customized high-performance detectors.

By strategically distributing optical power across lens groups, we placed high-power negative elements near the aperture stop to suppress Petzval and field curvature. Incorporation of advanced dispersion glasses enabled precise matching of chromatic dispersion, correcting both lateral and longitudinal chromatic aberrations and suppressing secondary spectra for APO performance. Joint optimization of the objective and tube lens provided additional design freedom, allowing fine aberration balancing and near diffraction-limited RMS wavefront error across the full field and multiple wavelengths. This approach yielded a water-immersion objective with an 8-mm FOV and 2-mm working distance, paired with a matched tube lens.

We employed real-time adaptive optics to correct peripheral aberrations within excitation spectral range at the field edges, a method experimentally validated through high-resolution imaging of dendritic spines. High-performance, large-aperture GaAsP PMTs were integrated into the design, with the objective optimized to make full use of the effective photocathode area, maximizing photon collection and minimizing noise level. This significantly enhanced the dynamic range of the imaging signal, particularly in deep tissue. The reason for this is that the PMT delivers an anode current that must be converted to voltage across a load or a transimpedance feedback resistor, which sets a fundamental trade-off between voltage gain, Johnson noise, and bandwidth. Lowering the resistance suppresses thermal noise and extends the bandwidth but reduces output voltage. Using high-output-current GaAsP PMTs resolves this trade-off by providing sufficiently large photocurrents to yield high output voltages even under low-resistance operation, thereby improving SNR and preserving temporal fidelity at short pixel dwell times ^13,49^.

The choice of PMT is an important design element for meso-scale two-photon microscopy, prioritizing signal linearity and collection geometry. A defining advantage is its high maximum anode output current, which establishes the broad dynamic range essential for quantitative calcium imaging. This capability is critical for resolving the high-amplitude fluorescence excursions associated with neuronal bursting, ensuring that these intense, rapid transients are digitized linearly without saturation or clipping. Furthermore, the expanded photocathode area minimizes vignetting by maximizing the capture of both highly scattered and ballistic photons. We do note the device’s lower quantum efficiency compared to to Multi-Pixel Photon Counters (SiPMs), which use Geiger-mode Avalanche Photodiodes. However, it is free from issues such as the reduction in response bandwidth caused by the dead time in Geiger mode.

Notably, in complex multiphoton optical systems, it is important to optimize dispersion effectively, as due to the broad spectral bandwidth (∼20 nm) associated with femtosecond pulses, material dispersion arising from optical glass components causes temporal pulse broadening. This lowers the temporal photon density, significantly reducing two-photon excitation efficiency at a fixed average power. We therefore performed GDD pre-compensation using the laser’s built-in grating module (ALCOR 920-4-XSight, Spark Lasers; tunable range: 0 to -60,000 fs). The compensation was tuned using image fluorescence intensity as a feedback variable. The optimal operating point was identified at a GDD value of -47,000 fs, where peak fluorescence intensity confirmed the restoration of the narrowest pulse width at the objective.

We conducted multi-scale imaging experiments on a variety of murine cortical structures, as well as large-scale neuronal ensemble calcium activity imaging, to demonstrate the system’s capability for high-resolution imaging across extensive fields and depths. To the best of our knowledge, this is currently the only two-photon microscope capable of simultaneously achieving ultra-wide-field imaging (50.27 mm^2^) together with cellular resolution (∼1 µm lateral), and deep-tissue penetration (up to 900 µm) in vivo at the multiple scales shown here.

Extended FOV multiphoton microscopy has been the subject of rapid advances over the last few years, with the introduction of a number of systems which have extended the state of the art, including 2P-RAM^10^, FASHIO-2PM^13^ and Diesel2p^12^. ULTRA exceeds the performance of these systems, especially in terms of FOV, ULTRA achieves an imaging area exceeding 50 mm² (Supplementary Fig. 10), compared to approximately 19.5 mm² for 2P-RAM, 9 mm² for FASHIO-2PM, and roughly 25 mm² for Diesel2p. Moreover, ULTRA further maintains the resolution capabilities of extended large-field two-photon microscopy for in vivo, deep-tissue imaging, achieving imaging depths of up to 900 μm in vivo, substantially extending the imageable volume achieved to 45.24 mm^3^ in the mouse brain.

Compared to these mainstream wide-field two-photon microscopes, our system exhibits relatively modest resolution, achieving a lateral resolution of 1.16 µm and an axial resolution of 11.33 µm (compared to 0.61 µm lateral and 4.52 µm axial for 2P-RAM; 1.61 µm lateral and 7.07 µm axial for FASHIO-2PM; and approximately 1 µm lateral and 8 µm axial for Diesel2p). Although numerically slightly lower than the resolutions of these other systems, a lateral resolution of around 1 µm is typically sufficient for most practical experimental scenarios. Conversely, there is considerable scope for improving axial resolution to enhance the system’s optical sectioning capability. However, improved axial resolution is a trade-off, as increased axial precision can adversely affect imaging stability, particularly during experiments involving awake and task-performing animals. Because a tighter point spread function (PSF) concentrates photon flux and thereby improves two-photon excitation efficiency, the optimal axial resolution must be selected in accordance with the demands of the experiment.

Due to the inherent performance constraints of available scanning components, our system does not significantly surpass other devices in terms of imaging speed. Given that even under maximum imaging conditions, ∼250 mW of under-objective laser power remained unused, sufficient power is left available to incorporate an additional scanning pathway (similar to the approach taken in Diesel2p^12^) potentially doubling the imaging speed without compromising the original imaging quality.

Lastly, our system has a working distance of 2 mm, shorter than the >3 mm provided by 2P-RAM, 4.5 mm by FASHIO-2PM, and 8 mm by Diesel2p. Given the greater capability of ULTRA for deep tissue imaging, extending the working distance would enhance usability in specific experimental contexts, particularly for imaging larger mammals, which possess thicker skulls and cortical layers.

Overall, the advances in ultra-wide-field multiphoton imaging capability reported here open up new avenues for neuroscientific research, providing unprecedented tools for exploring complex neural dynamics across diverse biological contexts. We expect that imaging the activity of tens of thousands of neurons distributed across multiple layers and the breadth of the mouse cortex during behaviour will be routinely achievable in the coming years, with current rapid progress suggesting that even greater sampling of brain circuit activity will soon be possible.

### Statistics and reproducibility

All imaging experiments were categorized based on specific experimental setups. For structural imaging of brain tissues: full-field neuronal somata imaging, n = 2 (Fig. 3a); wide-field dendritic spine imaging with deep-layer z-stacks, n = 2 (Fig. 3b); long-term dendritic spine tracking imaging, n = 3 (Fig. 3c); and large-scale vascular imaging across four independent regions, n = 2 (Fig. 3d). For cellular calcium activity imaging: single wide FOV (single hemisphere) superficial imaging, n = 3 (Figs. 4b and 5c); wide FOV superficial imaging spanning bilateral hemispheres, n = 4 (n = 1 for Fig. 4a; n = 3 for Fig. 5f); large-span superficial layer imaging across 4 independent regions (4 x 0.5mm x 0.5mm), n = 3 (Fig. 4d); and large-span deep-layer imaging across 4 independent regions (4 x 0.5mm x 0.5mm), n = 4 (n = 3 for Fig. 4g; n = 1 for Fig. 5h).

## Data availability

Data sufficient to reproduce the figures in this paper will be made available via DataDryad (datadryad.org) at the time of publication, with the link to be provided here upon acceptance. Raw imaging data from the studies reported here are available upon reasonable request from the corresponding author.

## Code availability

Analysis code will be provided upon reasonable request.

## Acknowledgements

We thank the kindness of Dr. Huanhuan Zeng, Haiyang Chen and Prof. Zhiheng Xu for providing transgenic mice. We thank Ann Go for advice on the presentation of behavioral detection, and Ann Go and Amanda Foust for helpful comments on an earlier version of this manuscript. This work was supported by National Key Research and Development Program (2024YFC3406602) to G.Y. ; The Major scientific research facility project of Jiangsu Province (BM2022010), National Key R&D Program of China (2023YFC2413100), The Basic Research Pilot Project of Suzhou (No. SJC2021021); The Youth Innovation Promotion Association, CAS (No. 2022328), and National Natural Science Foundation of China (32127801, 61705251) to H.J. & Z.-Q.Z, and CAS Special Program for Research Facility Improvement (292025000036), a CAS PIFI fellowship to S.R.S., and EPSRC EP/W024020/1 ; EPSRC EP/Y020316/1; Royal Society IEC\NSFC\242498; and Chan-Zuckerberg Initiative Collaborative Pairs Award CP-2-1-Schultz to S.R.S.

## Contributions

M.Y. motivated the project and worked with Z.-Q.Z., Y.G., S.S. and Y.T. conceived the study. M.Y. designed the study, metrics and benchmark pipeline and collected the datasets. H.J. coordinated the facility and study. M.Y. and Z.-Q.Z. built the microscope with air-floating platform. Yan Gong designed the optics and mechanical structure with S.L., H.Z., L.Z., and Z.Z.. Z.-Q.Z. and S.S. perform the AO correction. M.Y. performed the imaging experiments. M.Y., T.L., E.S., J.Y. and Y.L. analyzed the datasets. M.Y., Z.-Q.Z., G.Y. and S.S. wrote the paper with contribution from all authors. M.Y. led the project.

## Supplementary information

**Description of Additional Supplementary Files (video)**

**Supplementary Video 1**

**Supplementary Video 2**

**Supplementary Video 3**

**Supplementary Video 4**

**Supplementary Video 5**

**Supplementary Video 6**

**Supplementary table 1.** Summary of the resolution measurement across the whole FOV.

| A | X-axis |  | Y-axis |  | Z-axis |  |
| --- | --- | --- | --- | --- | --- | --- |
| Orientation | Mean | SD | Mean | SD | Mean | SD |
| Center | 1.02 | 0.04 | 1.06 | 0.02 | 10.76 | 0.25 |

| Ring B | X-axis |  | Y-axis |  | Z-axis |  |
| --- | --- | --- | --- | --- | --- | --- |
| Orientation | Mean | SD | Mean | SD | Mean | SD |
| 0° | 1.13 | 0.09 | 1.07 | 0.03 | 10.52 | 0.08 |
| 45° | 1.17 | 0.06 | 1.08 | 0.02 | 10.84 | 0.15 |
| 90° | 1.17 | 0.06 | 1.05 | 0.03 | 11.23 | 0.02 |
| 135° | 1.149 | 0.08 | 1.04 | 0.02 | 10.79 | 0.17 |
| 180° | 1.12 | 0.07 | 1.10 | 0.08 | 10.72 | 0.12 |
| 225° | 1.18 | 0.11 | 1.06 | 0.03 | 10.61 | 0.06 |
| 270° | 1.15 | 0.14 | 1.04 | 0.02 | 10.25 | 0.06 |
| 315° | 1.16 | 0.11 | 1.05 | 0.02 | 11.17 | 0.13 |

| Ring C | X-axis |  | Y-axis |  | Z-axis |  |
| --- | --- | --- | --- | --- | --- | --- |
| Orientation | Mean | SD | Mean | SD | Mean | SD |
| 0° | 1.11 | 0.10 | 1.09 | 0.01 | 10.65 | 0.15 |
| 45° | 1.17 | 0.13 | 1.08 | 0.03 | 11.75 | 0.12 |
| 90° | 1.28 | 0.05 | 1.03 | 0.03 | 10.77 | 0.11 |
| 135° | 1.22 | 0.07 | 1.06 | 0.03 | 10.39 | 0.13 |
| 180° | 1.19 | 0.11 | 1.09 | 0.77 | 10.13 | 0.19 |
| 225° | 1.18 | 0.09 | 1.06 | 0.02 | 9.93 | 0.17 |
| 270° | 1.23 | 0.10 | 1.05 | 0.03 | 10.79 | 0.17 |
| 315° | 1.17 | 0.06 | 1.14 | 0.08 | 10.53 | 0.20 |

| Ring D | X-axis |  | Y-axis |  | Z-axis |  |
| --- | --- | --- | --- | --- | --- | --- |
| Orientation | Mean | SD | Mean | SD | Mean | SD |
| 0° | 1.13 | 0.12 | 1.63 | 0.04 | 13.14 | 0.20 |
| 45° | 1.32 | 0.02 | 1.28 | 0.05 | 13.99 | 0.26 |
| 90° | 1.78 | 0.15 | 1.14 | 0.04 | 14.00 | 0.18 |
| 135° | 1.39 | 0.06 | 1.26 | 0.05 | 13.50 | 0.21 |
| 180° | 1.10 | 0.06 | 1.44 | 0.06 | 11.64 | 0.24 |
| 225° | 1.21 | 0.11 | 1.21 | 0.04 | 12.52 | 0.22 |
| 270° | 1.65 | 0.10 | 1.13 | 0.09 | 12.54 | 0.31 |
| 315° | 1.29 | 0.09 | 1.27 | 0.02 | 13.16 | 0.21 |

**Supplementary Fig. 1.**
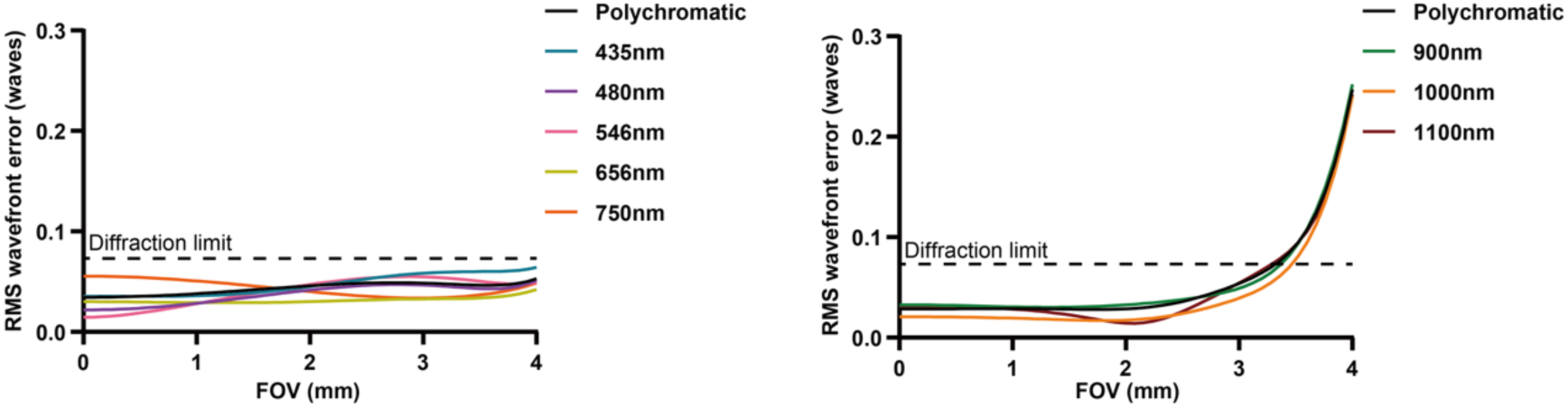
Simulated wavefront aberrations and diffraction-limited performance. **Left**, RMS wavefront error of the objective lens as a function of field of view (FOV) across visible and short-NIR wavelengths (435–750 nm). The design maintains diffraction-limited performance (RMS error < 0.07 waves, indicated by the dashed horizontal line) across the entire 4 mm FOV. **Right**, RMS wavefront error of the integrated optical system (objective and tube lens) for near-infrared wavelengths (900–1100 nm). The system remains diffraction-limited within 0–0.85 times of the full FOV (up to ∼3.4 mm), ensuring high-fidelity signal collection for deep-tissue imaging. These results validate the achromatic and wide-field capabilities of the custom-designed optics for multi-modal neural recording.

**Supplementary Fig. 2.**
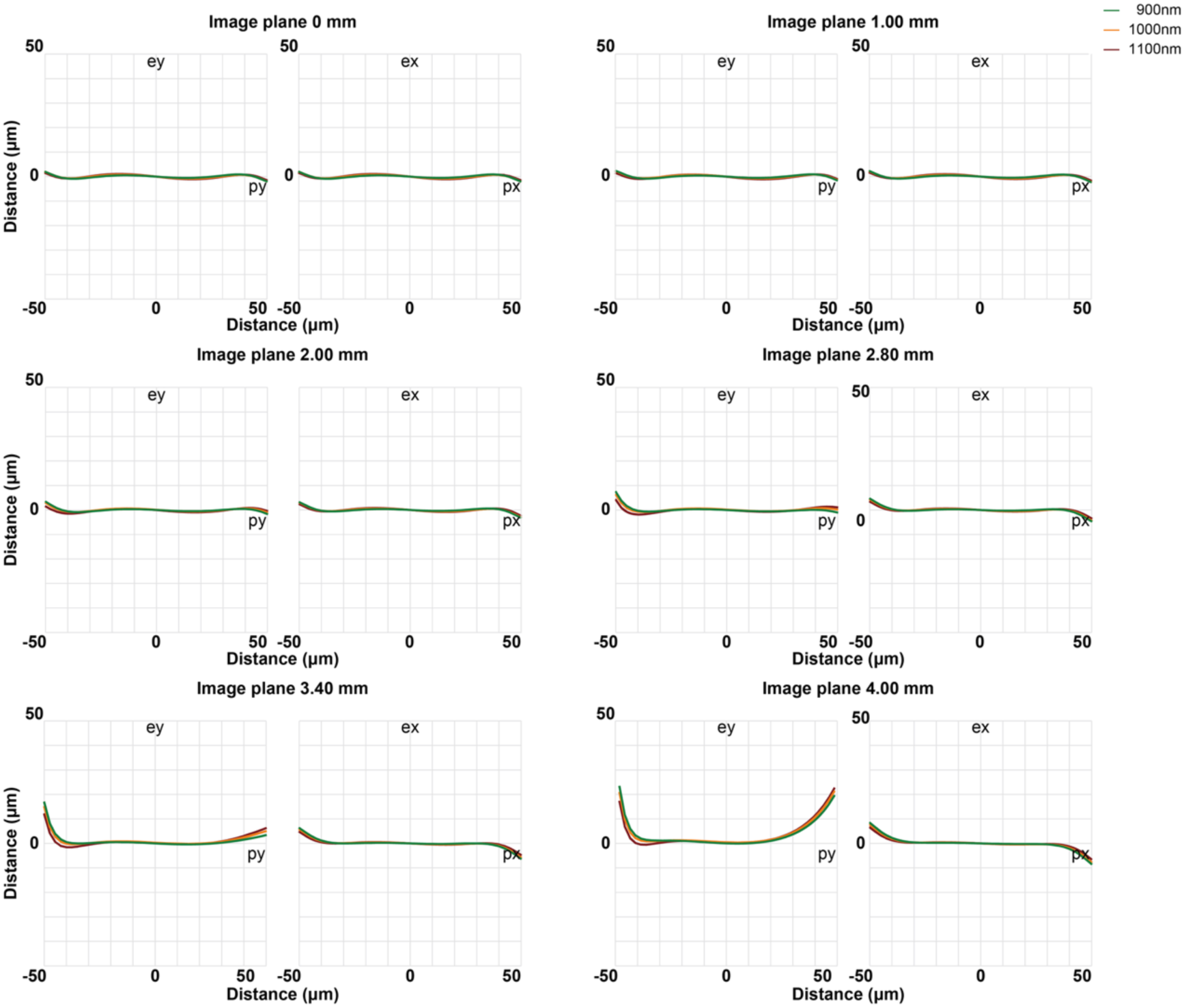
Optical aberration characterization. Simulated RMS wavefront error as a function of field of view (FOV) for the objective lens alone (435–750 nm) and the integrated system including the tube lens (900–1100 nm). The design achieves diffraction-limited performance across the entire 4 mm FOV for visible wavelengths and up to 0.85 times FOV (∼3.4 mm) for the NIR regime. Transverse ray aberration curves of the integrated system at representative field positions (0, 1.00, 2.00, 2.80, 3.40, and 4.00 mm) for NIR wavelengths (900, 1000, and 1100 nm). While the on-axis performance (0 mm) exhibits minimal residual spherical aberration, off-axis fields are primarily dominated by coma. Despite these aberrations, the system maintains high-fidelity imaging quality within the designated functional FOV.

**Supplementary Fig. 3.**
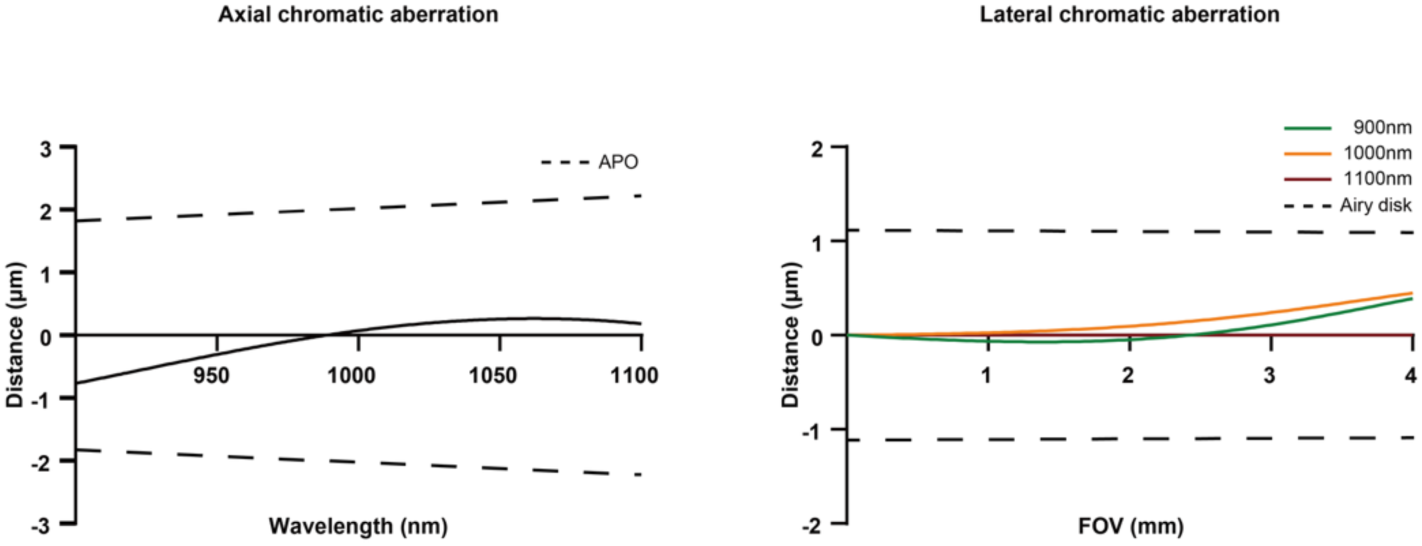
Chromatic aberration correction. Simulated spectral transmittance of the objective lens, showing >90% throughput from 435 to 1100 nm. **Left**, Axial chromatic aberration (longitudinal focal shift) of the objective-tube lens system from 900 to 1100 nm. The shift is calculated based on Zernike Z_4 = 0 and remains well within the APO tolerance limits (dashed lines). **Right**, Lateral chromatic aberration as a function of FOV for 900, 1000, and 1100 nm (referenced to 1100 nm). The lateral shifts across the entire FOV are contained within the Airy disk radius (dashed lines), ensuring spatial co-registration of multi-spectral NIR signals without computational alignment.

**Supplementary Fig. 4.**
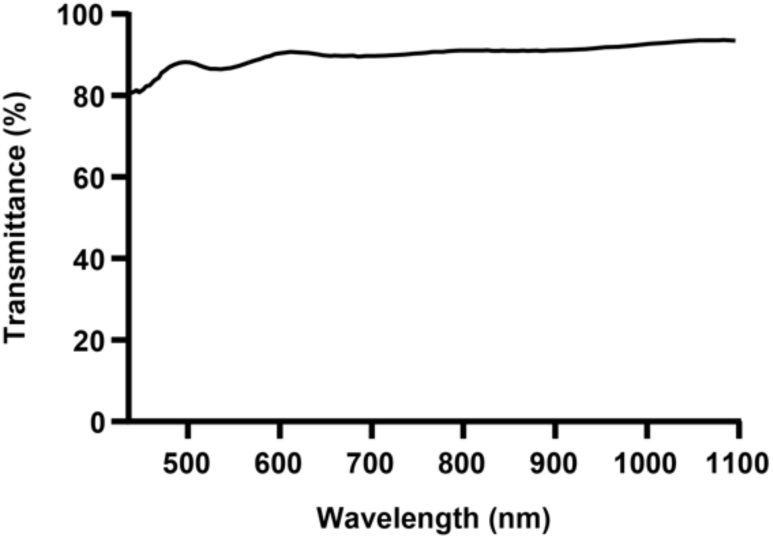
Simulated broadband spectral transmittance of the custom-designed objective. The simulated transmittance profile of the objective lens demonstrates high throughput (exceeding 90% across most regions) over a broad spectral range from 435 nm to 1100 nm. This bandwidth covers the entire visible spectrum and extends into the NIR regime, ensuring efficient photon collection for multi-modal imaging and deep-tissue applications.

**Supplementary Fig. 5.**
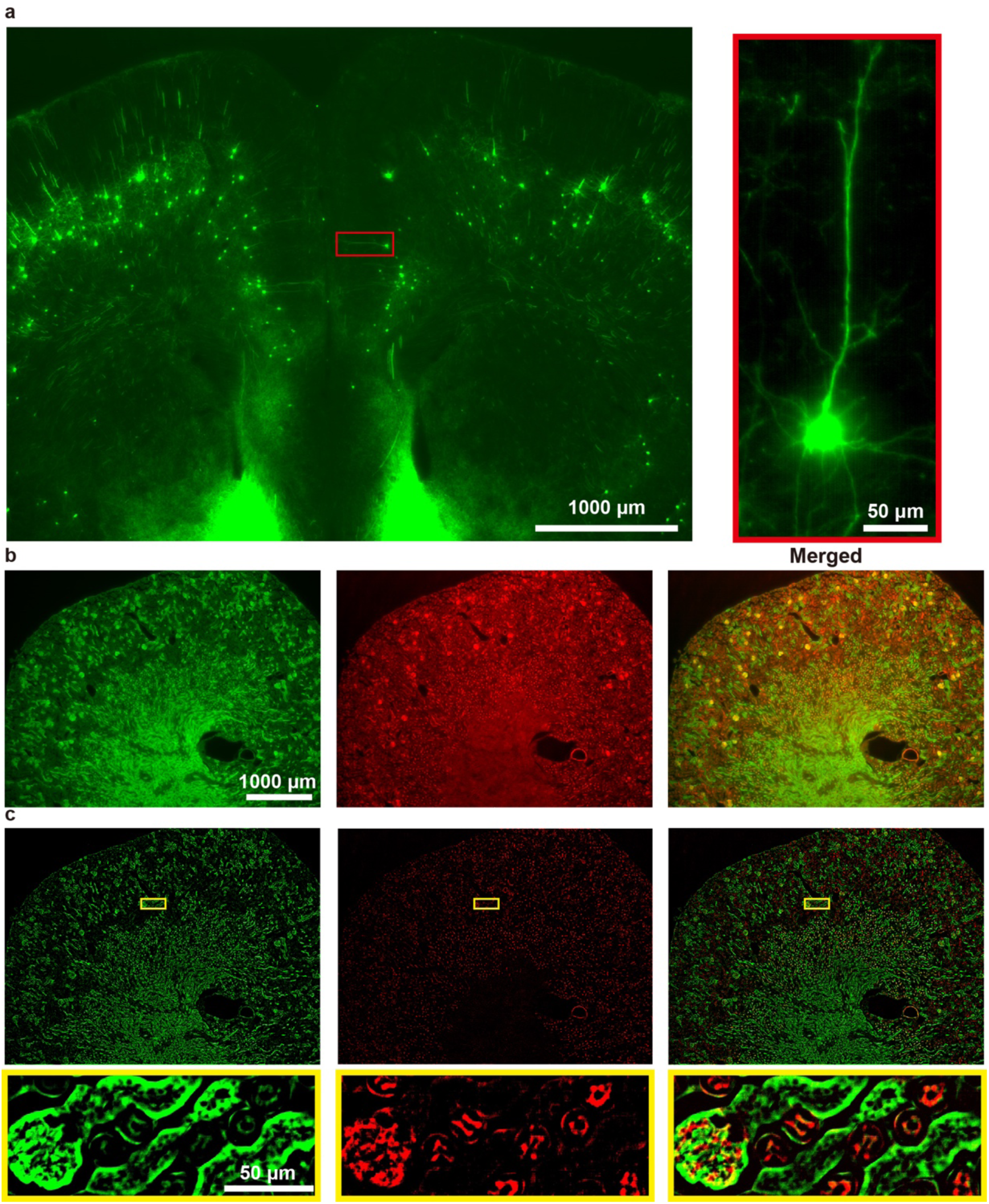
Wide-field fluorescence imaging of various tissue samples using ULTRA’s objective. **a**, Wide-field fluorescence imaging of brain slice of Thy1-GFP transgenic mice (thickness 50 μm) pixel size 14192 x 10640, FOV 4807 μm x 3604 μm. **b**, Dual-color wide-field fluorescence imaging of mouse kidney (FluoCells™ Prepared Slide #3 (Cat. No. F24630)), elements of the glomeruli and convoluted tubules stained in green-fluorescent lectin and filamentous actin prevalent stained in red-fluorescent phalloidin. **c**, Speckle illumination dual-color wide-field fluorescence imaging of same sample in **c** enhancing optical sectioning.

**Supplementary Fig. 5.**
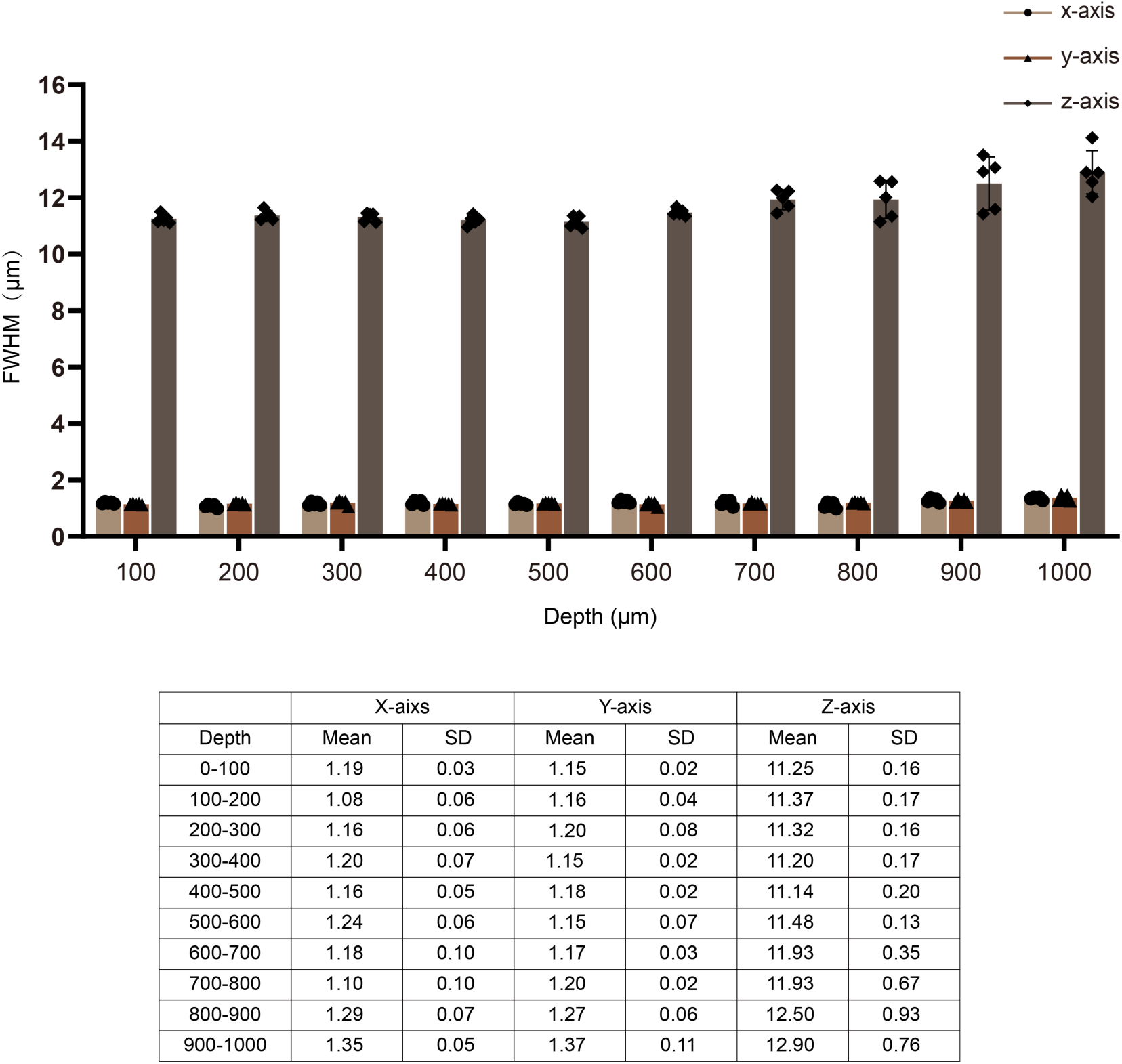
Resolution measurement in depth. 0.5-μm fluorescent microspheres embedded in a thick 0.75% agarose gel, were imaged at the center of the FOV under the objectives and 10 100-μm z-stack were acquired from o to 1000 μm deep in the sample (n=5). The FWHM of the Gaussian fits for measurements (mean ± SD) indicate both lateral and axial resolutions maintain good consistency as imaging depth increases, relative to the resolutions measured at the sample surface **(all measurements are in μm)**.

**Supplementary Fig. 7.**
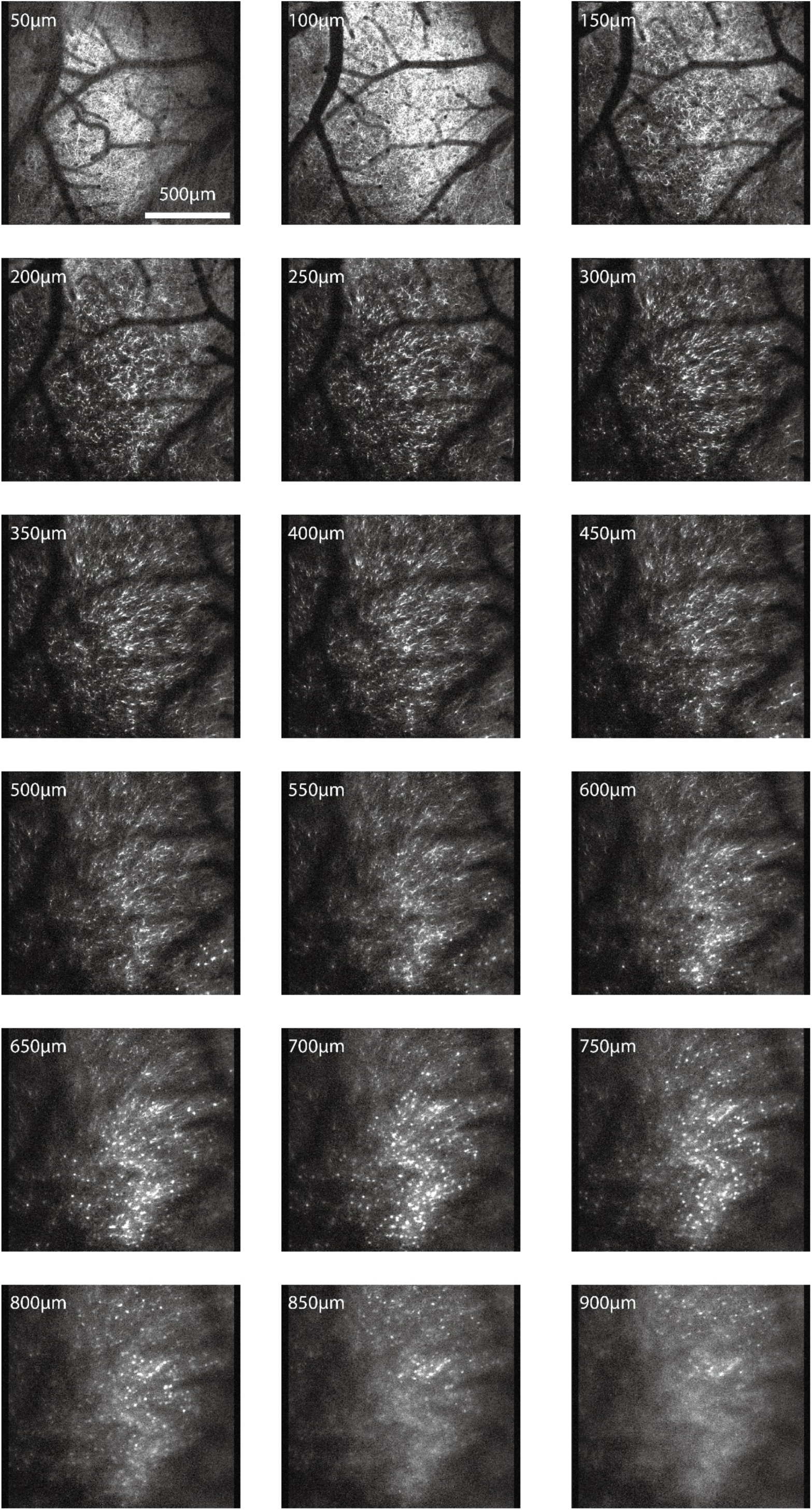
ULTRA z-stack imaging of dendrites ana somata up to 900 μm (50 μm per step) shown in Fig. 3b right.

**Supplementary Fig. 8.**
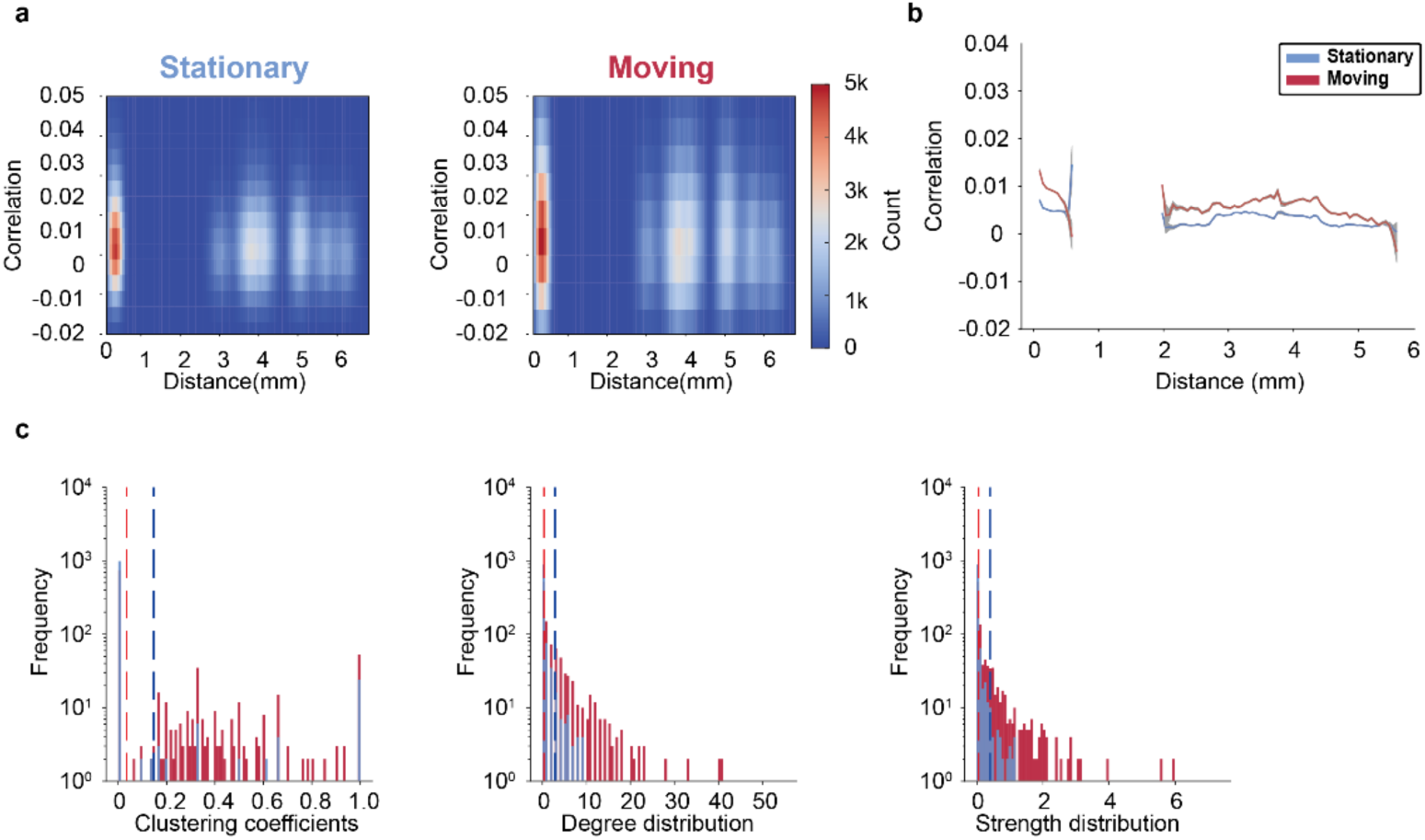
Functional connectivity assessments on neuronal ensembles across 4 independent regions. **a**, Distribution of correlation as a function of distance between neurons between states. **b**, Correlation as a function of distance between neurons between states. **c**, Similar analysis in Fig. 5d with mean value (dashed line).

**Supplementary Fig 9.**
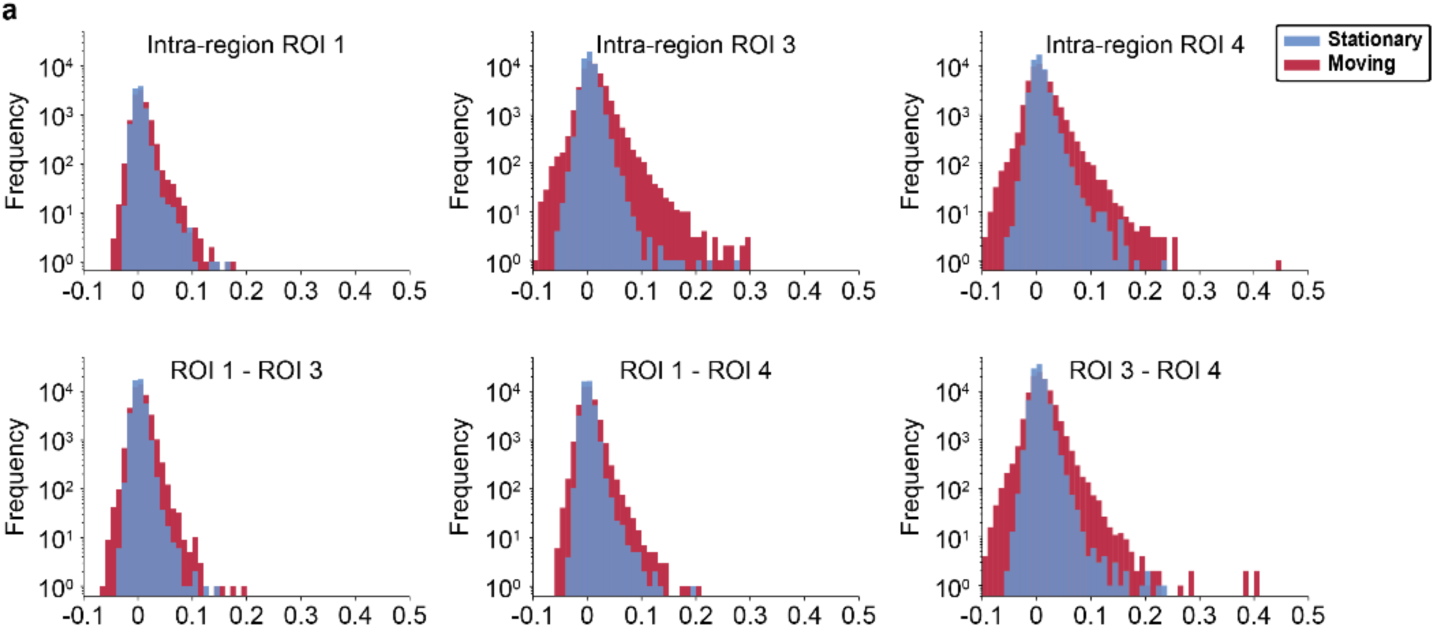
Correlation distribution of intra-region and inter-region between states.

**Supplementary Fig. 10.**
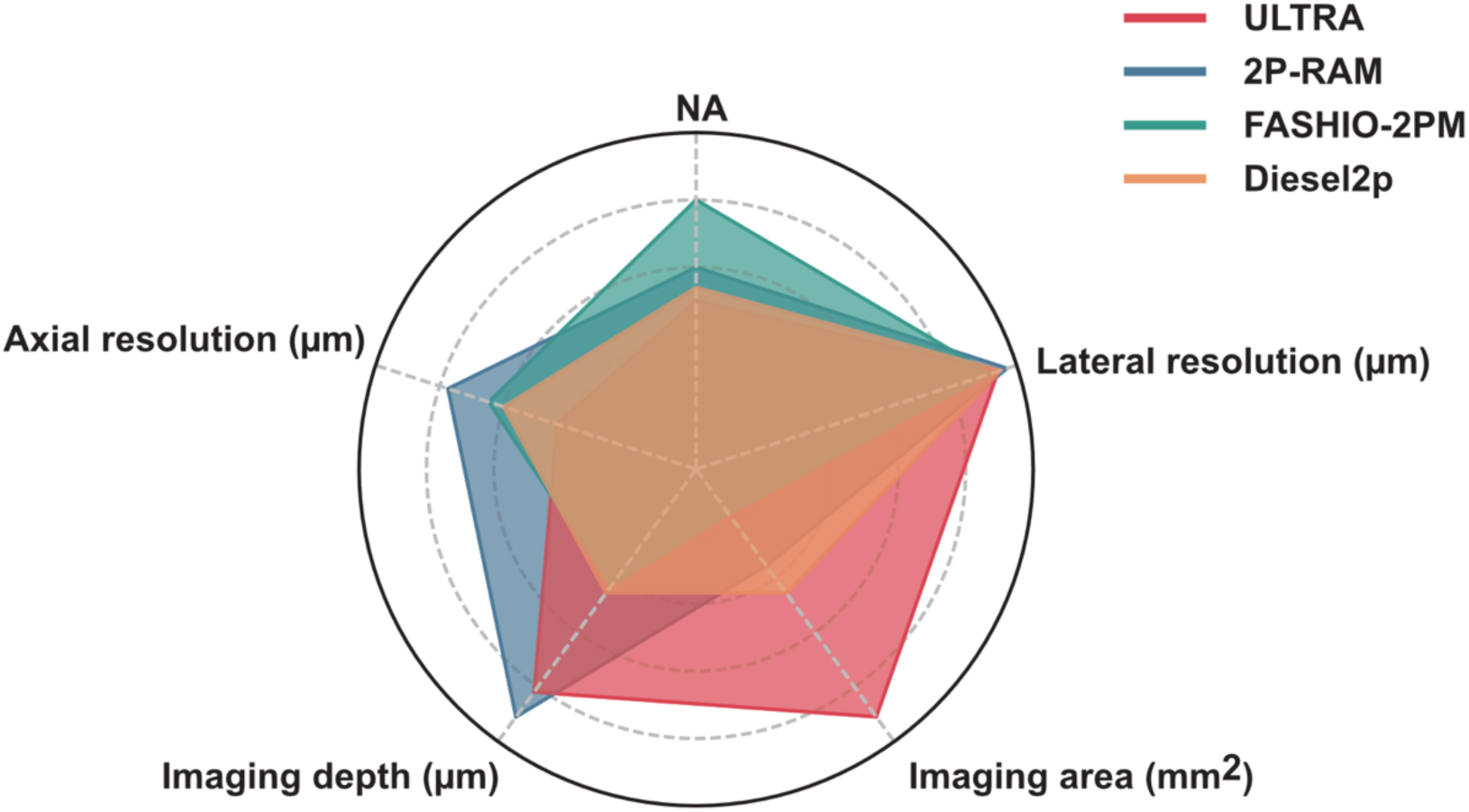
Comparison of ULTRA with several other recently developed two-photon mesoscopes. Performance comparison of ULTRA with state-of-the-art multi-photon systems. The radar chart summarizes five key optical performance metrics (all the values are ranged from the center outwards): Numerical Aperture (NA, range 0-1), Lateral resolution (μm, range 20-0), Imaging area (mm^2^, range 0-55), Imaging depth (μm, range 0-1100), and Axial resolution (μm, range 20-0). ULTRA (red) demonstrates a significant advantage in imaging area while maintaining high NA and spatial resolution, effectively achieving a higher space-bandwidth product (SBP) compared to 2P-RAM (blue), FASHIO-2PM (green), and Diesel2p (orange).

